# Atypical brain asymmetry in autism – a candidate for clinically meaningful stratification

**DOI:** 10.1101/2020.03.24.000349

**Authors:** Dorothea L. Floris, Thomas Wolfers, Mariam Zabihi, Nathalie E. Holz, Marcel P. Zwiers, Tony Charman, Julian Tillmann, Christine Ecker, Flavio Dell’Acqua, Tobias Banaschewski, Carolin Moessnang, Simon Baron-Cohen, Rosemary Holt, Sarah Durston, Eva Loth, Declan Murphy, Andre Marquand, Jan K. Buitelaar, Christian F. Beckmann, the EU-AIMS LEAP group

## Abstract

**Background:** Autism Spectrum Disorder (henceforth ‘autism’) is a highly heterogeneous neurodevelopmental condition with few effective treatments for core and associated features. To make progress we need to both identify and validate neural markers that help to parse heterogeneity to tailor therapies to specific neurobiological profiles. Atypical hemispheric lateralization is a stable feature across studies in autism, however its potential of lateralization as a neural stratification marker has not been widely examined.

**Methods:** In order to dissect heterogeneity in lateralization in autism, we used the large EU-AIMS Longitudinal European Autism Project dataset comprising 352 individuals with autism and 233 neurotypical (NT) controls as well as a replication dataset from ABIDE (513 autism, 691 NT) using a promising approach that moves beyond mean-group comparisons. We derived grey matter voxelwise laterality values for each subject and modelled individual deviations from the normative pattern of brain laterality across age using normative modeling.

**Results:** Results showed that individuals with autism had highly individualized patterns of both extreme right- and leftward deviations, particularly in language-, motor- and visuospatial regions, associated with symptom severity. Language delay (LD) explained most variance in extreme rightward patterns, whereas core autism symptom severity explained most variance in extreme leftward patterns. Follow-up analyses showed that a stepwise pattern emerged with individuals with autism with LD showing more pronounced rightward deviations than autism individuals without LD.

**Conclusion:** Our analyses corroborate the need for novel (dimensional) approaches to delineate the heterogeneous neuroanatomy in autism, and indicate atypical lateralization may constitute a neurophenotype for clinically meaningful stratification in autism.

## Introduction

Autism spectrum disorder (autism) is a neurodevelopmental condition characterized by social-communicative deficits, restricted, repetitive behaviors and sensory abnormalities (1). One of the key characteristics of autism is its great phenotypic and biological heterogeneity (2,3). Particularly in the neuroimaging literature, results are mixed and inconsistent and have been attributed to this heterogeneity in the autism population. To provide a more coherent picture of the complex neuropathology of autism, it is critical to tackle its heterogeneity and identify the neural markers that are consistent across studies. Such markers can provide clinically relevant stratification of individuals with autism.

When it comes to identifying a consistently implicated neural feature in autism, a large body of literature converges towards a disruption of hemispheric specialization - one of the most fundamental biological properties of our brains (4,5). This basic organizational principle describes that the two hemispheres differ in their functional specialization and exhibit pronounced structural asymmetries (6,7). Functional specialization involves leftward lateralization of language and motor skills and rightward lateralization of spatial perceptual abilities (8). It is also evident in grey matter (GM) asymmetries with frontal opercular and temporal perisylvian regions, and hippocampus exhibiting leftward asymmetries, whereas thalamus and posterior parietal cortex showing rightward asymmetries (9,10).

Individuals with autism exhibit impairments in left hemisphere skills such as social-communication, language and motor-related symptoms, whilst appearing relatively intact in right hemisphere functions such as visuospatial skills (11). This lateralized pattern of deficits and strengths in autism has given rise to theories trying to reconcile its complex clinical profile with atypical structural hemispheric specialization (11). In the largest autism cohort to date comprising over 3000 subjects individuals with autism presented with widespread leftward cortical reductions (12). Smaller studies are in line with this showing either a reduction or even reversal of typical leftward asymmetries in language- and motor-related regions (13–20). Thus, atypical lateralization in autism is among the most replicated findings with moderate to high effect sizes (21) in the otherwise heterogeneous neuroimaging literature. However, inconsistencies in results remain regarding regional specificity and direction of observed patterns. Such inconsistencies are usually attributed to sample heterogeneity and delineating heterogeneity still remains one of the central tasks in autism research.

The relationship between heterogeneity and brain asymmetry in autism may be further co-dependent on age, sex, handedness, different symptom profiles and comorbidities such as attention-deficit hyper-activity disorder (ADHD). *Age*: Lateralization in neurotypicals becomes more pronounced through age-related maturational processes (22). This typical trajectory is disrupted in individuals with autism showing increasing reversed rightward lateralization (23). *Sex*: NT males are usually more strongly lateralized, while NT females have a more symmetric distribution pattern (24). How sex affects atypical asymmetry in autism is unknown. *Handedness*: Left-handed individuals have a higher chance of a different organization in the brain than right-handed individuals. Individuals with autism exhibit elevated rates of non-right-handedness, which has been attributed to atypical specialization in the brain (25). *Language delay (LD):* Individuals with autism and early LD show more pronounced deviations from typical asymmetry than those without LD (15,26). *ADHD*: Some conditions that are also a common comorbidity in autism, such as ADHD, are also associated with atypical lateralization (27).

Despite increased recognition of heterogeneity in autism (28), there is little effort to address such challenges methodologically. One first example towards quantifying biological variation at the individual level in autism was recently demonstrated (29) using a novel normative modeling method (30,31). Similar to the use of growth charts in paediatric medicine, normative modelling aims to place each individual with respect to centiles of variation in the population and thereby facilitates a move away from classical case-control analyses that ignore individual differences. Applying normative modelling to cortical thickness estimates, Zabihi et al., 2018 (29) showed that individuals with autism exhibit highly individualized atypicalities of cortical development.

This study was designed to address heterogeneity in autism with regard to age, sex and core and co-occurring symptoms in the context of brain lateralization using novel individualized analyses: 1) we transcend classical case-control analysis and address inter-individual variation by applying normative modeling (30,32). 2) We aim to identify laterality-related subtypes by considering co-occurring clinical symptoms in autism such as language development. Through capturing variation at the individual level in combination with addressing different sources of heterogeneity and using a consistent imaging feature in autism, our work provides a step towards precision neuroscience in autism.

## Methods and Materials

### Participants

Participants were part of the EU-AIMS and AIMS-2-TRIALS Longitudinal European Autism Project (LEAP) (33,34) cohort - the largest European multi-center initiative aimed at identifying biomarkers in autism. Participants underwent comprehensive clinical, cognitive and MRI assessment at one of six collaborating sites (Figure S1). All participants with ASD had an existing clinical diagnosis of autism. For details on participants, study design and exclusion criteria, see Supplemental Information (SI) and (34). The final sample comprised 352 individuals with autism (259 males and 93 females), and 233 NT controls (154 males and 79 females) between 6 and 30 years. For details on demographic information, see Table 1.

### Clinical and cognitive measures

IQ was assessed using the Wechsler Abbreviated Scales of Intelligence. The Autism Diagnostic Observation Schedule (ADOS-G (35)) measured clinical core symptoms of autism. The Autism Diagnostic Interview-Revised (ADI-R) (36) was used to measure parent-rated autistic symptoms and LD, which was defined as having onset of first words later than 24 months and/or first phrases later than 33 months. ADHD symptoms were assessed with the DSM-5 ADHD rating scale. Handedness was assessed with the short version of the Edinburgh Handedness Inventory (37). For further details on all cognitive measures, see SI and (33).

### Image Preprocessing

For MRI data acquisition parameters, see SI and Table S1. Structural T1-weighted images were preprocessed according to a validated laterality pipeline (15,38,39) using the CAT12 toolbox (http://www.neuro.unijena.de/cat) (see Figure S2). All original images were segmented and affine registered to a symmetric tissue probability map before being reflected across the cerebral midline (x◻=◻0). All segmented reflected and original (non-reflected) grey matter (GM) maps were then used to generate a symmetrical study-specific template via a flexible, high-dimensional nonlinear diffeomorphic registration algorithm (DARTEL) (40). They were next registered to the symmetrical study-specific template as per standard DARTEL procedures. An intensity modulation step was included to retain voxel-wise information on local volume (39). The final resulting images were modulated, warped, reflected (I_ref_) and non-reflected (I_nref_) GM intensity images. A laterality index (LI) was calculated at each voxel using the following formula: LI = 2(I_nref_−I_ref_)/(I_nref_ + I_ref_). Positive values in the right hemisphere of the LI image indicate rightward asymmetry, and negative values in the right hemisphere of the LI image indicate leftward asymmetry. Values of LI in the left hemisphere have identical magnitude, but opposite sign and were therefore excluded from further analyses. LI images were smoothed with a 4-mm FWHM isotropic Gaussian kernel.

### Normative Modelling

The normative modeling method has been described in detail previously (29,30,41–43). In summary, we estimated a normative brain aging model at each GM laterality voxel by using Gaussian process regression (GPR) (44), a Bayesian non-parametric interpolation method that yields coherent measures of predictive confidence in addition to point estimates (for details see SI). With this method, we could predict both the expected regional GM asymmetry changes and the associated predictive uncertainty for each individual allowing us to quantify the voxelwise deviation of GM asymmetry from the NT range across the entire brain.

First, we trained a GPR model at each voxel on the NT cohort using age, sex and site as covariates to predict GM asymmetry resulting in a developmental model of GM asymmetry in NT individuals. To avoid over-fitting, assess generalizability and determine whether NT individuals fall within the normative range, we used 10-fold cross-validation in NT individuals before retraining the model in the entire sample to make predictions in individuals with autism (see SI).

We generated normative probability maps (NPM), which quantify the deviation of each participant from the normative model for GM asymmetry at each voxel. These subject-specific Z-score images provide a statistical estimate of how much each individual’s true laterality value differs from the predicted laterality value with reference to the NT pattern at each voxel given the participants age, sex and site. NPMs were thresholded at an absolute value of Z>|2.6| (30,42). Based on this fixed threshold, we defined extreme rightward-lateralized (positive) and extreme leftward-lateralized (negative) deviations for each participant.

All extreme deviations per subject were summarized into (log-transformed) scores representing the percentage of extreme rightward and extreme leftward deviations per subject in relation to the total number of intracerebral voxels. These percentage scores were compared calculating a GLM including diagnosis and sex as the regressors of interest, and age as covariate. To compare our normative modeling against a conventional group-mean difference analysis we ran Permutation Analysis of Linear Models (PALM) on the LI images examining the sex-by-diagnosis(-by-age) interaction with site as covariate.

### Spatial characterization of deviations

We applied two strategies to spatially characterize the extreme rightward and leftward deviations: 1) We generated spatial overlap maps for individuals in the same diagnostic and gender group by summing up the number of extreme deviations in each voxel for each subject. These were then divided by the total number of subjects (multiplied by 100) to represent the percentage of extreme right- and leftward deviations per brain voxel. 2) We extracted extreme rightward and leftward deviations within structurally (Harvard-Oxford atlas (HOA (45))), and functionally (neurosynth (http://neurosynth.org, accessed June 2019) (46)) defined regions of interest (ROIs) (see SI). For latter, we used the search terms ‘language’, ‘motor’ (left-lateralized), ‘visuospatial’, ‘attention’ (right-lateralized) and ‘monitoring’, ‘mentalizing’ (no lateral bias). Results were FDR-corrected. Effect sizes were computed using Cohen’s *d*. For robustness and replicability analyses, see SI.

### Relative contributions of different cognitive and behavioral measures

We ran relative importance analyses within individuals with autism including variables related to lateralization such as LD, ADHD, handedness, sex and symptom severity (CSS). This was done in a sample with reduced sample size (N=305) without missing values on these variables. We used averaging over orderings (47) to rank the relative contribution of highly correlated regressors to the linear regression model. The variable showing strongest contribution to explained variance was followed-up with additional analyses.

### Symptom Associations

An individual-level atypicality score was estimated for each individual through extreme value statistics by computing the 10% trimmed mean of the 1% top deviations (29). We then computed one-tailed, FDR-corrected Pearson’s correlations between these individual atypicality scores and the ADI-R and ADOS-2 symptom severity scores.

### Robustness and Replicability

To assess robustness, we estimated a separate normative model without including site as a covariate, but removing site effects using ComBat (48). For the sake of comparability, we also applied FDR-correction (49) to the NPMs. We also included full-scale intelligence (FIQ) and handedness as nuisance covariates. To test whether the effects were robust against the influence of intellectual disability (ID), we reran analyses excluding individuals with FIQ <70. Finally, as individuals with autism with and without LD were not matched on certain demographic variables, we additionally created a sub-sample matched for age and symptom severity and re-ran second-level statistical analyses to assess robustness of results. To assess replicability, we selected a sample from the publically available Autism Brain Imaging Data Exchange (ABIDE) I and II (50,51) to examine extreme right- and leftward deviations in an independent datasets. For details, see the SI and Table S2.

## Results

The classic PALM analysis to assess mean group differences did not yield any significant results. Figure 1 depicts the spatial representation of the voxel-wise normative model in NT males in the largest site KCL (N=42). Results were comparable for NT females (Figure S3) and when using ComBat to address site effects (Figures S4a and S4b). The forward model for all other sites by sex is shown in Figure S5. Both in NT males and females leftward shifts in GM asymmetry were mainly evident in Heschl’s gyrus, planum temporale, parietal and central operculum, angular gyrus (AG), supramarginal gyrus (SMG), pars triangularis, postcentral gyrus, precentral gyrus, superior and middle frontal gyrus, lateral occipital cortex (LOC) and cerebellum. Rightward shifts in GM asymmetry mainly occurred in middle and inferior temporal gyrus, posterior SMG, AG, superior parietal lobule, precuneus, anterior cingulate cortex, supracalcarine cortex, lingual gyrus, occipital pole, and caudate.

**Figure 1.**
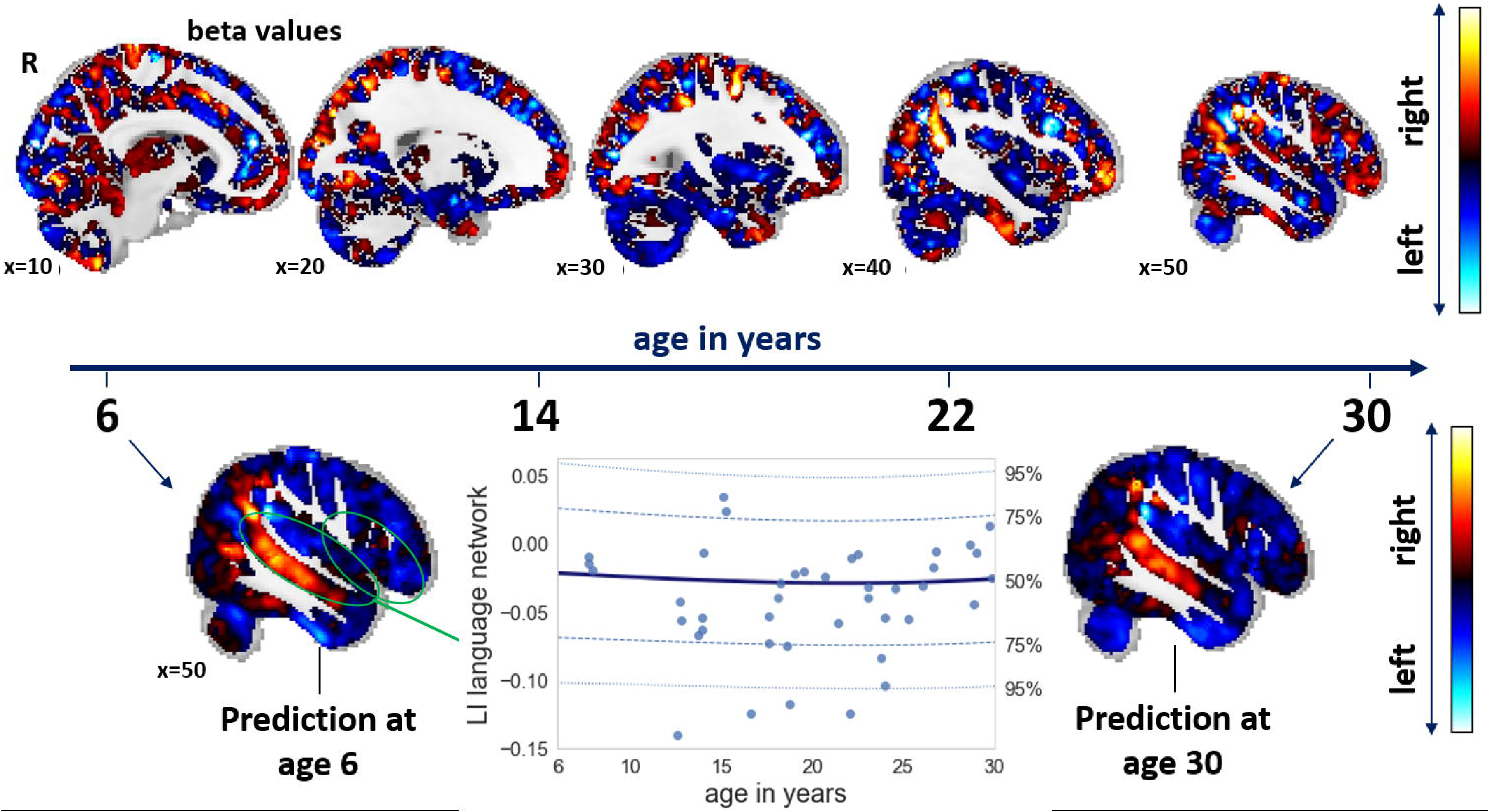
Normative developmental changes for GM laterality in neurotypical males. Figure 1 captures the spatial representation of the voxelwise normative model in neurotypical (NT) males from the KCL site. The upper panel shows the beta values of laterality change across 6 to 30 years of age. The lower panel shows the actual prediction of GM laterality at ages 6 and 30. Blue colors indicate a shift towards leftward asymmetry, whereas red colors indicate a shift towards rightward asymmetry. The regression line depicts the predicted laterality values extracted from the language network based on neurosynth between 6 and 30 years of age along with centiles of confidence. These are based on the normative model maps thresholded with positive Rho-maps. Blue dots are the true values for NT males in KCL.

The accuracy of the normative model was evaluated using the correlation between the true and the predicted voxel values (Rho) generated under 10-fold cross-validation. NT Rho maps were restricted to positive values and further analyses were conducted on normative model maps thresholded with these. For unthresholded Rho maps along with root mean square error maps, see Figures S6 and S7.

### Characterization of extreme deviations

Overall, males and females with autism showed both significantly more extreme rightward (*F*_*(1)*_=12.5, *p*<0.001, *d*=0.31) and leftward deviations (*F*_*(1)*_=12.0, *p*<0.001, *d*=0.29) compared to NT males and females (Figure 2). There were no sex differences (right: *F*_*(1)*_=0.2, *p*=0.67, *d*=0.06; left: *F*_*(1)*_=0.2, *p*=0.67, *d*=0.06) nor sex-by-diagnosis interactions (right: *F*_*(1)*_=2.2, *p*=0.14, *d*=0.26; left: *F*_*(1)*_=0.5, *p*=0.47, *d*=0.11). Controlling the FDR at the individual level of each NPM led to identical conclusions (see SI and Figure S8). Overall individuals with autism showed a trend towards more extreme right-than leftward deviations (χ^*2*^=3.59, *p*=0.05). Further, there was a significant positive correlation between extreme right- and leftward deviations (*r*=0.41, *p*<0.001). For rightward deviations, females with autism showed the highest overlap in parahippocampal gyrus, putamen and amygdala, while males with autism mostly in the middle temporal gyrus and hippocampus (Figure 2a). For leftward deviations, females with autism showed highest overlap in orbitofrontal cortex, frontal pole and postcentral gyrus, while males with autism mostly in LOC, temporal pole and thalamus (Figure 2b). On average females with autism had higher overlapping deviating regions than males with autism (right: χ^*2*^=667, *p*<0.001; left: χ^*2*^=277, *p*<0.001).

**Figure 2.**
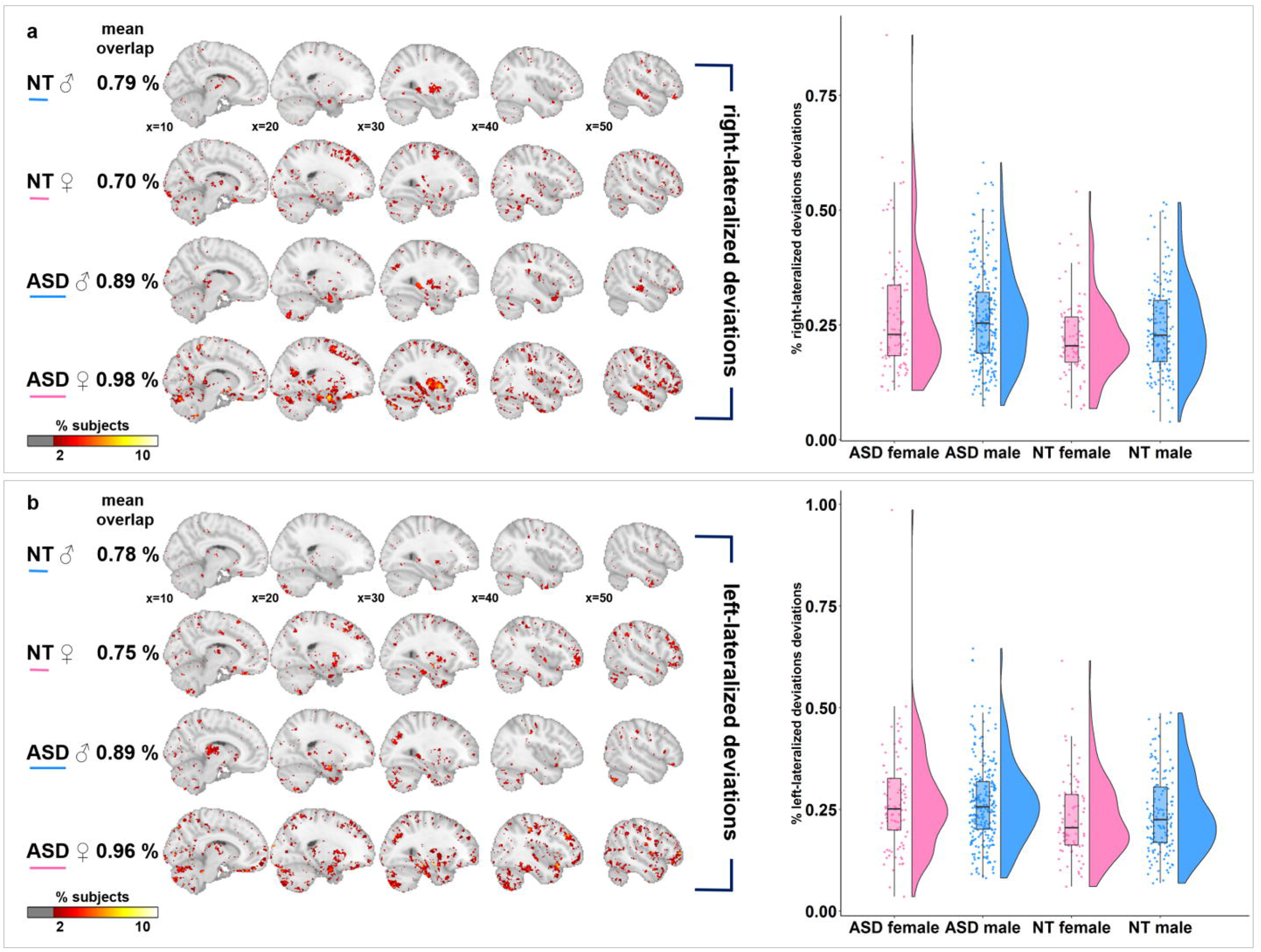
Characterization of extreme laterality deviations. Abbreviations: ASD=autism spectrum disorder, NT=neurotypicals. The upper panel characterizes extreme rightward deviations (a), the lower panel extreme leftward deviations (b). The left panels show the percentage of extreme right- and leftward deviations from the normative model at each brain locus for each diagnostic group and gender separately. We depict loci where at least 2% of the subjects show overlaps. The violin plots on the right show the extreme deviations for each individual within each diagnostic and gender group. On average, individuals with autism show more extreme deviations than NT controls for both right- and leftward deviations.

When considering deviations by structural ROIs, we found a significant main effect of diagnosis in the frontal (*F*_*(1)*_=15.7, *p*<0.001, *d*=0.35) and central operculum (*F*_*(1)*_=14.9, *p*<0.001, *d*=0.34) with individuals with autism showing more extreme rightward deviations, and in the superior lateral occipital cortex (*F*_*(1)*_=14.4, *p*<0.001, *d*=0.32) with individuals with autism showing more extreme leftward deviations. Further, there was a significant sex-by-diagnosis interaction in the temporal occipital fusiform cortex (TOFC) (*F*_*(1)*_=19.2, *p*<0.001, *d*=0.79) with females with autism having more extreme rightward deviations. When considering deviations by functional ROIs, individuals with autism showed extreme rightward deviations in the motor network (*F*_*(1)*_=9.8, *p*=0.001, *d*=0.27), whereas more extreme leftward deviations in the visuospatial network (*F*_*(1)*_=11.6, *p*<0.001, *d*=0.3). Results were robust across sensitivity analyse and replicated in the large ABIDE sample (see SI).

### Relative contributions

Results revealed that LD had relatively the largest importance for explaining extreme rightward deviations (LD: 62.1% > handedness: 17.3% > sex: 10.9% > ADHD: 9.9% > CSS: 0.5%), while symptom severity for extreme leftward deviations (CSS: 40.4% > LD: 34.1% > sex: 23.6% > ADHD: 12.3% > handedness: 2.3%) (Figure 3). LD was further followed up showing a significant main effect for rightward deviations (*F*_*(2)*_=10.5, *p*<0.001). Specifically, individuals with autism and LD were different from both individuals with autism without LD (*t*_(213)_=2.5, *p*=0.01, *d*=0.32) and NT individuals (*t*_(152)_=4.6, *p*<0.001, *d*=0.58), while individuals with autism without LD were not different from NT individuals (*t*_(256)_=1.9, *p*=0.06, *d*=0.21). This stepwise pattern was overall more pronounced in males than in females with autism (Figure 4 and Figure S9).

**Figure 3.**
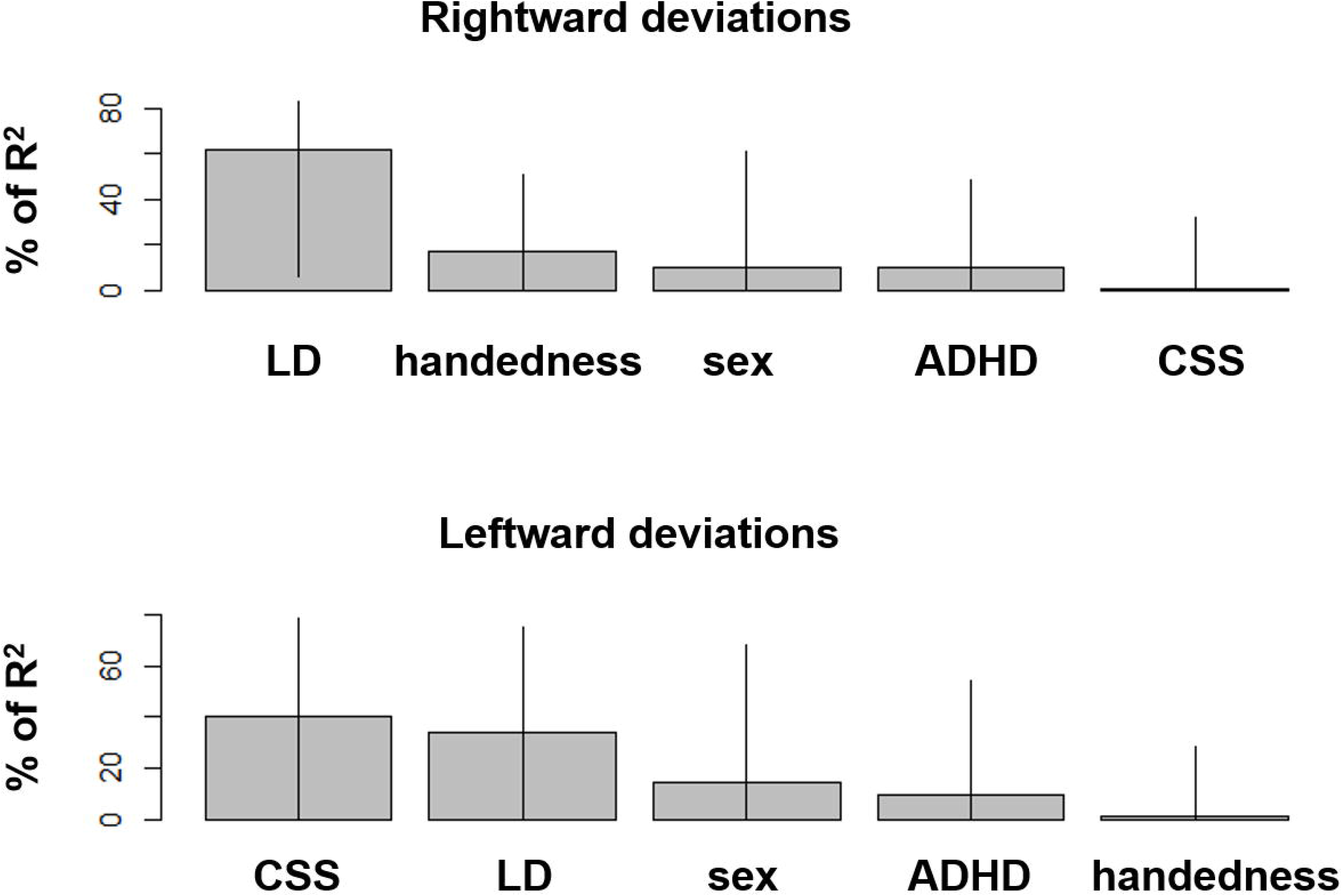
Relative importance of extreme laterality deviations. Abbreviations: LD=language delay, ADHD=attention-deficit hyper-activity disorder, CSS=calibrated severity scores on the ADOS-2. Based on the R package ‘relaimpo’ (https://cran.r-project.org/web/packages/relaimpo/relaimpo.pdf), LD has the largest contribution to R^2^ for extreme rightward deviations, whereas symptom severity has the largest contribution to R^2^ for extreme leftward deviations.

**Figure 4.**
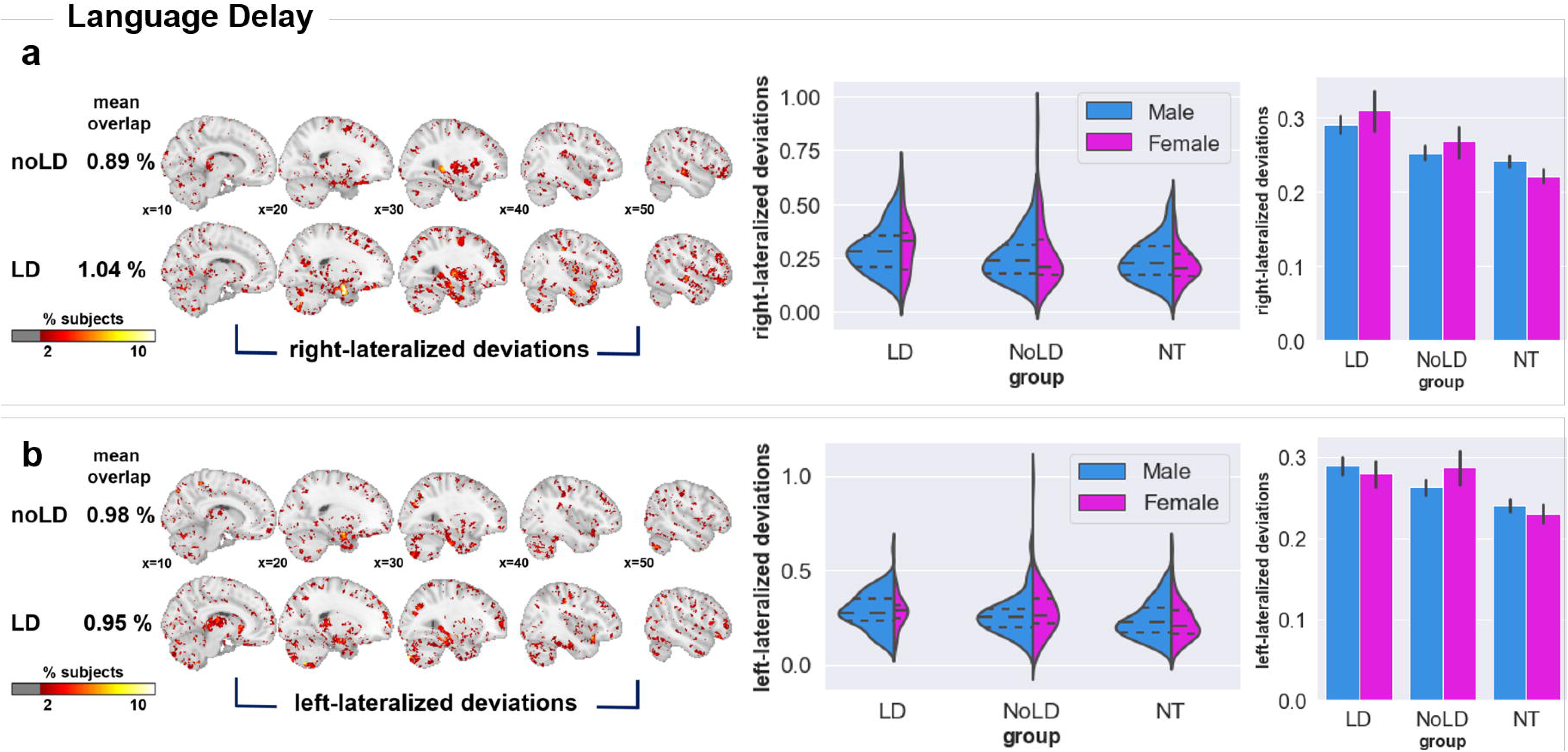
Extreme laterality deviations as a function of language delay. Abbreviations: LD=individuals with autism with language delay, noLD=individuals with autism without language delay, NT=neurotypicals. The upper panel characterizes extreme rightward deviations (a), the lower panel extreme leftward deviations (b). The left panels show the percentage of extreme right-and leftward deviations from the normative model at each brain locus for each diagnostic group and gender separately. We depict loci where at least 2% of the subjects show overlaps. The violin and bar plots on the right show that individuals with autism with LD show more extreme rightward deviations than the other two groups.

Results by language-related ROIs showed a main effect of LD for extreme rightward deviations in the frontal operculum (*F*_*(2)*_=10.6, *p*<0.001), the central operculum (*F*_*(2)*_=14.9, *p*<0.001) and the motor network (*F*_*(2)*_=6.9, *p*=0.001). Rightward deviations in the language network were significant (*F*_*(2)*_=4.1, *p*=0.01), but did not survive FDR-correction. For all, there was a stepwise pattern, with individuals with autism and LD showing more pronounced deviations. For details, see SI.

### Symptom Associations

Across the whole brain, there were significant associations between extreme rightward deviations and ADI-R communication scores (*r*=0.14, *p*=0.004). No correlations survived FDR-correction in individuals with autism with and without LD. In males with autism, there were significant positive correlations between rightward deviations and core autism symptoms (ADI-R social: *r*=0.18, *p*=0.002; ADI-R communication: *r*=0.19, *p*=0.001). In females with autism, as well as in males and females with autism with and without LD, no results survived FDR-correction.

### Replicability

In line with the results of the EU-AIMS LEAP sample, males and females with autism of the combined ABIDE dataset showed both significantly more extreme rightward (*F*_*(1)*_=7.7, *p*=0.006, *d*=0.14) and leftward deviations (*F*_*(1)*_=17.1, *p*<0.001, *d*=0.24) compared to NT individuals (see Figures S10a-b). As in EU-AIMS LEAP, there were no sex differences (right: *F*_*(1)*_=1.1, *p*=0.3, *d*=0.14; left: *F*_*(1)*_=2.9, *p*=0.13, *d*=0.19) and no sex-by-diagnosis interactions (right: *F*_*(1)*_=3.6, *p*=0.06, *d*=0.26; left: *F*_*(1)*_=3.4, *p*=0.06, *d*=0.26). When considering deviations by functional ROIs significant in the primary analysis, individuals with autism also showed extreme rightward deviations in the motor network (*F*_*(1)*_=5.2, *p*=0.02, *d*=0.21), and extreme leftward deviations in the visuospatial network (*F*_*(1)*_=4.0, *p*=0.05, *d*=0.12). There were no significant group differences in the frontal operculum (*F*_*(1)*_=0.2, *p*=0.6, *d*=0.02). For further details, see the SI and Figures 10a-b.

## Discussion

In this study, we mapped extreme deviations in structural asymmetry at the individual level in comparison to a normative model of laterality development. A further aim was to explore laterality as a stratification marker in autism in a large and deeply phenotyped cohort and an equally large replication cohort. We found highly individualized patterns of both right- and leftward asymmetry deviations in males and females with autism. In contrast, when using classical case-control analyses, we did not detect any significant group differences, emphasizing the need to move beyond group averages to capture an accurate representation of the phenotype at the individual level. Similarly, a recent study addressing heterogeneity dimensionally points out that traditional case-control analyses yield smaller effects and miss atypicalities detected with a dimensional subtyping approach (52). By deriving statistical inferences at the individual level, we demonstrated that individuals with autism show a different, individualized pattern that would otherwise be missed when focusing on a common neurobiological signature. In fact, spatial overlap of extreme deviations across subjects was minimal in both males and females with autism (Figure 2 and Figure S10) corroborating the high degree of heterogeneity across subjects.

Moreover, extreme right- and leftward deviations were highly correlated with each other, which suggests that biological factors acting on atypical laterality shifts might not be biased towards one side in the same individual in autism. It thus also appears that a disruption in establishing the typical laterality pattern is not confined to one hemisphere, but there are inter-related alterations in both. In line with several previous reports (13,15,53), our results also show that individuals with autism have slightly more extreme rightward than leftward deviations. The biological underpinning are unknown, but likely include both genetic (54), prenatal endocrine (55) and environmental factors. For instance, Geschwind et al., (2002) (56) reported that left frontal and extreme rightward deviations regions are under less genetic control as compared to the right hemisphere and therefore are more susceptible to environmental influences during neurodevelopment. Whether prenatal androgens and related early immune activation (55,57) contribute to atypical hemispheric development in autism, remains to be established.

Extreme rightward deviations were most pronounced in the motor network and frontal operculum and extreme leftward deviations in the visuospatial network. Accordingly, rightward shifts in asymmetry in language related regions, particularly in Broca’s area, are frequently reported in autism (16,17,58). However, atypical lateralization of motor and visuospatial performance is underexplored in autism. Accumulating evidence suggests the important role of motor-related asymmetries in the neurobiology of autism (5,13,59,60). Despite mostly intact visuospatial performance in autism, atypical activation patterns have also been reported in individuals with autism while performing visuospatial tasks (61). More specific cognitive measures are needed to establish the functional relevance of these alterations.

Males and females with autism showed a similar degree of extreme deviations (Figure 2). Specifically, in the TOFC, females with autism showed stronger rightward deviations, while males with autism and NT females both show fewer rightward deviations than NT males. This ‘occipital face area’ is strongly right-lateralized in NT individuals (62). Individuals with autism exhibit atypical face processing strategies and these are more pronounced in females with autism (63). Being one of the most reported impairments in social cognition in autism, face processing and related atypical, sex-differential lateralization awaits further exploration.

When considering the spatial extent of deviations, females with autism show on average greater overlap across differently implicated regions than males with autism (Figure 2 and Figure S10). This suggests that females with autism constitute a less heterogeneous group than males with autism who show less pronounced overlap of focal atypicalities in laterality. Also, when considering males and females separately, atypical asymmetry was only associated with more social-communicative symptoms in males with autism, but not females with autism, suggesting that phenotypically similar manifestations of atypical lateralization appear to have differential cognitive implications for males and females with autism.

We found that both extreme left- and rightward deviations were associated with LD, but only rightward deviations differentiated individuals with autism with and without LD from each other. This is in line with prior reports (15,26). Two structural studies further showed differential morphological alterations in individuals with autism with different developmental language profiles (64) and a loss of leftward asymmetry in the arcuate fasciculus in non-verbal children with autism (65). The degree of atypical rightward lateralization may thus constitute a biological marker of different etiological subgroups with different language profiles in autism. Evidence suggests onset of language before the age of two years (66), and the level of language at the ages five and six years (67), predict functional outcome later in life in autism. Developing stratification markers based on LD will thus have important clinical implications for early prognosis, individualized diagnoses and early language-based interventions in individuals at risk.

### Strengths and Limitations

We present analyses in a large-scale, deeply phenotyped and prospectively harmonized dataset. Results present robust across several analyses addressing potential confounds and are overall also observable in the large-scale, yet not harmonized ABIDE. Replicability and robustness of findings are important prerequisites as a first step towards establishing neural biomarkers.

Nevertheless, our findings should be considered in light of some limitations. 1) Despite the large dataset, when dividing individuals by diagnostic group, sex, LD and site, sub-samples decrease substantially in size. While our Bayesian GPR model adapts to the availability of data (i.e. gives more conservative estimates when data density decreases (31)), the model would yield more precise estimates with larger sample sizes. 2) Large datasets come at the expense of additional confounds associated with site. Here, we addressed this both by including site in the model and by running an alternative method for controlling batch effects. Our results are robust, however, residual site effects cannot fully be excluded. 3) The question arises whether there are also experience-dependent influences on cortical asymmetry. Longitudinal analyses are needed to pinpoint the onset and trajectory of atypical development in subgroups on the autism spectrum. 4) The normative modeling approach is suitable for detecting extreme deviations from a normative pattern. However, an alternative hypothesis that individuals with autism might lack specialization of either hemisphere (59) which would be expressed in reduced laterality might be less detected with the current approach.

## Conclusion

We estimated a normative model of GM voxelwise asymmetry based on a large NT cohort and applied this to a large, deeply phenotyped and heterogeneous autism sample. Our results confirm that atypical asymmetry is a core feature of the autism neurophenotype, shows highly individualized patterns across individuals and is differentially related to different symptom profiles such as language delay. Further exploration of such associations has the potential to yield clinically relevant stratification markers needed for precision medicine.

## Acknowledgements

We thank all participants and their families for participating in this study. We also gratefully acknowledge the contributions of all members of the EU-AIMS LEAP group: Jumana Ahmad, Sara Ambrosino, Bonnie Auyeung, Tobias Banaschewski, Simon Baron-Cohen, Sarah Baumeister, Christian F. Beckmann, Sven Bölte, Thomas Bourgeron, Carsten Bours, Michael Brammer, Daniel Brandeis, Claudia Brogna, Yvette de Bruijn, Jan K. Buitelaar, Bhismadev Chakrabarti, Tony Charman, Ineke Cornelissen, Daisy Crawley, Flavio Dell’Acqua, Guillaume Dumas, Sarah Durston, Christine Ecker, Jessica Faulkner, Vincent Frouin, Pilar Garcés, David Goyard, Lindsay Ham, Hannah Hayward, Joerg Hipp, Rosemary Holt, Mark H. Johnson, Emily J.H. Jones, Prantik Kundu, Meng-Chuan Lai, Xavier Liogier D’ardhuy, Michael V. Lombardo, Eva Loth, David J. Lythgoe, René Mandl, Andre Marquand, Luke Mason, Maarten Mennes, Andreas Meyer-Lindenberg, Carolin Moessnang, Nico Mueller, Declan G.M. Murphy, Bethany Oakley, Laurence O’Dwyer, Marianne Oldehinkel, Bob Oranje, Gahan Pandina, Antonio M. Persico, Barbara Ruggeri, Amber Ruigrok, Jessica Sabet, Roberto Sacco, Antonia San José Cáceres, Emily Simonoff, Will Spooren, Julian Tillmann, Roberto Toro, Heike Tost, Jack Waldman, Steve C.R. Williams, Caroline Wooldridge, and Marcel P. Zwiers. This project has received funding from the Innovative Medicines Initiative 2 Joint Undertaking under grant agreement No 115300 (for EU-AIMS) and No 777394 (for AIMS-2-TRIALS). This Joint Undertaking receives support from the European Union’s Horizon 2020 research and innovation programme and EFPIA and AUTISM SPEAKS, Autistica, SFARI. This work was also supported by the Netherlands Organization for Scientific Research through Vidi grants (Grant No. 864.12.003 [to CFB] and Grant No. 016.156.415 [to AFM]); from the FP7 (Grant Nos. 602805) (AGGRESSOTYPE) (to JKB), 603016 (MATRICS), and 278948 (TACTICS); and from the European Community’s Horizon 2020 Programme (H2020/2014-2020) (Grant Nos. 643051 [MiND] and 642996 (BRAINVIEW). This work received funding from the Wellcome Trust UK Strategic Award (Award No. 098369/Z/12/Z) and from the National Institute for Health Research Maudsley Biomedical Research Centre (to DM). TW gratefully acknowledges grant support from the Niels Stensen Fellowship. NEH gratefully acknowledges grant support from the German Research Foundation (grant numbers DFG HO 5674/2-1, GRK2350/1) and the Olympia Morata Program of the University of Heidelberg. SBC was funded by the Autism Research Trust, the Wellcome Trust, the Templeton World Charitable Foundation, and the NIHR Biomedical Research Centre in Cambridge, during the period of this work.

---

**Figure S1 – Distribution of subjects across acquisition sites**

Abbreviations: ASD=autism spectrum disorder, NT=neurotypicals. The bar plot shows the distribution of subjects by diagnostic group and sex across the five acquisition sites. Cambridge: autism males=34, autism females=16, NT males=19, NT females=9; KCL= autism males=103, autism females=33, NT males=42, NT females=25; Mannheim: autism males=22, autism females=6, NT males=25, NT females=10; Nijmegen: autism males=72, autism females=27, NT males=40, NT females=23; Utrecht: autism males=28, autism females=11, NT males=28, NT females=12.

**Figure S2 – Summary of GM VBM laterality pre-processing pipeline**

Abbreviations: GM=grey matter, VBM=voxel-based morphometry, DARTEL=Diffeomorphic Anatomical Registration using Exponentiated Lie algebra (40), LI=laterality index, I_nref_ =non-reflected GM images, I_ref_= reflected GM images. Figure depicts the pre-processing pipeline for structural T1-weighted images. Note that the standard VBM pipeline is adjusted to meet the needs for laterality analyses, i.e., using symmetric registration.

**Figure S3 – Normative developmental changes for GM laterality in neurotypical females**

Supplementary Figure 3 captures the spatial representation of the voxelwise normative model in neurotypical (NT) females from the KCL site. The upper panel shows the beta values of laterality change across 6 to 30 years of age. The lower panel shows the actual prediction of GM laterality at ages 6 and 30. Blue colors indicate a shift towards leftward asymmetry, whereas red colors indicate a shift towards rightward asymmetry. The regression line depicts the predicted laterality values extracted from the language network based on neurosynth between 6 and 30 years of age along with centiles of confidence. These are based on the normative model maps thresholded with positive Rho-maps. Blue dots are the true values for NT females in KCL.

**Figure S4a – Normative developmental changes for GM laterality in neurotypical males using ComBat**

Supplementary Figure 4a captures the spatial representation of the voxelwise normative model in neurotypical (NT) males across all sites. The upper panel shows the beta values of laterality change across 6 to 30 years of age. The lower panel shows the actual prediction of GM laterality at ages 6 and 30. Blue colors indicate a shift towards leftward asymmetry, whereas red colors indicate a shift towards rightward asymmetry.

The regression line depicts the predicted laterality values extracted from the language network based on neurosynth between 6 and 30 years of age along with centiles of confidence. These are based on the normative model maps thresholded with positive Rho-maps. Blue dots are the true values for NT males across all sites.

**Figure S4b – Normative developmental changes for GM laterality in neurotypical females using ComBat**

Supplementary Figure 4b captures the spatial representation of the voxelwise normative model in neurotypical (NT) females across all sites. The upper panel shows the beta values of laterality change across 6 to 30 years of age. The lower panel shows the actual prediction of GM laterality at ages 6 and 30.

Blue colors indicate a shift towards leftward asymmetry, whereas red colors indicate a shift towards rightward asymmetry. The regression line depicts the predicted laterality values extracted from the language network based on neurosynth between 6 and 30 years of age along with centiles of confidence. These are based on the normative model maps thresholded with positive Rho-maps. Blue dots are the true values for NT females across all sites.

**Figure S5 – Normative model by sex and site in NT individuals for the language network**

The regression lines depict the predicted laterality values extracted from the language network based on neurosynth between 6 and 30 years of age along with centiles of confidence. These are based on the normative model maps thresholded with positive Rho-maps. Blue dots are the true values for NT males on the left and NT females on the right categorized by sites.

**Figure S6 – Mean model accuracy 1**

The figure shows the correlation (Rho) between true and predicted grey matter laterality values in NT controls and autism individuals.

**Figure S7 – Mean model accuracy 2**

The figure shows the root mean square error of true and predicted mean of grey laterality values in NT controls and autism individuals.

**Figure S8 – Characterization of extreme laterality deviations using FDR-correction**

Abbreviations: ASD=autism spectrum disorder, NT=neurotypicals, LD=individuals with autism with language delay, noLD=individuals with autism without language delay. The figure depicts the replication of results when using FDR-correction as an alternative thresholding method. The upper panel characterizes extreme rightward deviations (a), the lower panel extreme leftward deviations (b). The left panels show the percentage of extreme right- and leftward deviations from the normative model at each brain locus for each diagnostic group and gender separately. We depict loci where at least 2% of the subjects show overlaps. The violin plots in the middle show the extreme deviations for each individual within each diagnostic and gender group. On average, individuals with autism show more extreme deviations than NT controls for both right- and leftward deviations. The barplots on the right show that individuals with autism with LD have more extreme-rightward deviations.

**Figure S9 – Extreme laterality deviations as a function of language delay in matched subsamples**

Abbreviations: LD=individuals with autism with language delay, noLD=individuals with autism without language delay, NT=neurotypicals. Replicated results are shown in a sample where individuals with autism with and without LD are matched for age and symptom severity. The upper panel characterizes extreme rightward deviations (a), the lower panel extreme leftward deviations (b). The violin and bar plots on the right show that individuals with autism with LD show more extreme rightward deviations than the other two groups.

**Figure S10 – Overlap between ABIDE and EU AIMS**

Abbreviations: ASD=autism spectrum disorder, NT=neurotypicals, ABIDE=Autism Brain Imaging Data Exchange. The figure depicts the replication of results in an independent dataset. The upper panel characterizes extreme rightward deviations (a), the lower panel extreme leftward deviations (b). The left panels show the percentage of extreme right- and leftward deviations from the normative model at each brain locus for each diagnostic group and gender separately where ABIDE and EU-AIMS LEAP show overlap. The violin plots on the right show the extreme deviations for each individual within each diagnostic and gender group in the ABIDE dataset. On average, individuals with autism show more extreme deviations than NT controls for both right- and leftward deviations

**Table 1 – Demographic and clinical characterization of the LEAP sample**

Abbreviations: ASD=Autism Spectrum Disorder; NT=neurotypical, M=males, F=females, ADI-R= Autism Diagnostic Interview-Revised, ADOS=Autism Diagnostic Observation Schedule, RRB=restricted, repetitive behavior, CSS=calibrated severity score, ADHD= attention-deficit hyper-activity disorder, LD=language delay.

**Table S1 – Summary of acquisition parameters across sites**

**Table S2 – Demographic and clinical characterization of the ABIDE sample**

Abbreviations: ASD=Autism Spectrum Disorder; NT=neurotypical, M=males, F=females, ADI-R= Autism Diagnostic Interview-Revised, ADOS=Autism Diagnostic Observation Schedule, RRB=restricted, repetitive behavior, CSS=calibrated severity score ^a^Full-Scale IQ information was available for 384 ASD M, 87 ASD F, 435 NT M and 197 NT F. ^b^Verbal IQ information was available for 330 ASD M, 76 ASD F, 382 NT M and 159 NT F. ^c^Performance IQ information was available for 335 ASD M, 78 ASD F, 398 NT M and 173 NT F. ^d^ADI-R Social Scores were available for 341 ASD M and 74 ASD F. ^e^ADI-R Communication and RRB Scores were available for 342 ASD M and 74 ASD F. ^f^ADOS-Gotham Social-Affect Scores were available for 132 ASD M and 28 ASD F. ^g^ADOS-Gotham RRB Scores were available for 134 ASD M and 29 ASD F. ^h^ADOS-Gotham Severity Scores were available for 134 ASD M and 30 ASD F.

