## Supplemental Material for "Atypical brain asymmetry in autism – a candidate for clinically meaningful stratification"

**Supplementary Information**

**Participants, study design and exclusion criteria**

All participants with autism had an existing clinical diagnosis of autism according to DSM-IV (1), DSM-IV-TR (2), DSM-5 (3) or ICD-10 (4) criteria. Participants underwent comprehensive clinical, cognitive and MRI assessment at one of six collaborating sites: the Institute of Psychiatry, Psychology and [Neuroscience](https://www.sciencedirect.com/topics/neuroscience/neurosciences), King’s College London (KCL), London, United Kingdom; Autism Research Centre at the University of Cambridge, Cambridge, United Kingdom; Radboud University Nijmegen Medical Centre, Nijmegen, the Netherlands; University Medical Centre Utrecht, Utrecht, the Netherlands; Central Institute of Mental Health, Mannheim, Germany; and University Campus Bio-Medico, Rome, Italy. For a distribution of participants by diagnostic group and sex across the sites, see Figure S1. Exclusion criteria included the presence of any MRI contraindications (e.g. metal implants, braces, claustrophobia) or failure to give informed written consent to MRI scanning, as well as significant hearing or visual impairments not corrected by glasses or hearing aids. In addition, we excluded participants with missing T1-weighted MRI scans, low image quality (i.e., structural brain abnormality, excessive head motion, insufficient coverage), and failed image preprocessing. Due to low number of participants from the Rome site after quality control, we restricted analyses to the five remaining sites. Neurotypical controls (NT) scoring positively for attention-deficit hyperactivity disorder (ADHD) as assessed by the DSM-5 ADHD rating scale were also excluded. The study was approved by the local ethical committees of the participating centers and written informed consent was obtained from all participants or their legal guardians.

**Clinical and cognitive measures**

General intellectual abilities were assessed using the Wechsler Abbreviated Scales of Intelligence-Second Edition (WASI-II (5)), or if unavailable the Wechsler Intelligence Scale for Children-III/IV (WISC-III/IV (6,7)) for children or Wechsler Adult Intelligence Scale for Adults-III/IV (WAIS-III/IV (8,9)) for adults. Standardized estimates of verbal IQ (VIQ), performance IQ (PIQ), and full-scale IQ (FIQ) were derived using IQ norms with mean=100 and SD=±15.

The Autism Diagnostic Observation Schedule (ADOS-G (10)) was used to measure the impact of current, clinically observed core symptoms of autism. Based on ADOS-2 algorithm totals, we report ADOS-2 Calibrated Severity Score (CSS) for ‘Social Affect’ indexing social-communication difficulties and ‘RRBs’ indexing restricted and repetitive behaviour. CSS Total serves an overall indicator of ASD severity. The CSS ranges from 1 to 10, with higher scores indicating more severe ASD symptom severity.

The Autism Diagnostic Interview-Revised (ADI-R) (11) is a structured parent interview completed by parents or caregivers of participants with autism. Algorithm scores were derived from current and historical symptom information for the domains of Reciprocal Social Interaction, Communication, and Restricted, Repetitive and Stereotyped Behaviors and Interests. The ADI‐R also assessed history of language development. Language delay was defined as having onset of first words later than 24 months and/or having onset of first phrases later than 33 months. ADHD symptoms were assessed with the DSM-5 ADHD rating scale, covering both inattention and hyperactivity/impulsivity symptoms based on either self-or parent-report (12). Self-report scores were only used when parent-report scores were unavailable (N=83). A categorical variable was computed based on the DSM-5 criteria (i.e., at least five positive responses in children and six in adults on either or both scales). Handedness was assessed with the short version of the Edinburgh Handedness Inventory (13). Scores ranged between +500 (right-handed) and -500 (left-handed). A categorical variable was computed comprising right-handed (+500 to +150), ambidextrous (-149 to +149) and left-handed (-150 to -500). For detailed information on clinical characteristics, see (14).

**MRI Data Acquisition**

All participants were scanned with a contemporary MRI scanner operating at 3T at 5 different sites (University of Cambridge: Siemens Verio; King’s College London: GE Medical Systems Discovery MR 750; Mannheim University: Siemens TimTrio; Radboud University: Siemens Skyra; Rome University: GE Medical Systems Signa HDxTt; Utrecht University: Philips Medical Systems Achieva/Ingenia CX). High-resolution structural T1-weighted volumetric images were acquired with full head coverage, at 1.2-mm thickness with 1.2x1.2-mm in-plane resolution. For all other scanning parameters, please see Table S2. Consistent image quality was ensured by a semi-automated quality control procedure.

**Gaussian process regression**

Gaussian process regression (GPR) (15) was used to estimate separate normative models of grey matter (GM) laterality at each voxel. Other methods are also suited to this purpose (e.g. Bayesian polynomial regression), GPR provides superior estimation of the mean and the ability to map the variation across the cohort through centiles of predictive confidence. For a full treatment of Gaussian processes see (16,17).

A Gaussian process (GP) specifies a distribution over functions, such that any finite

number of elements has a joint Gaussian distribution. They are excellent tools for Bayesian regression: given a dataset specified by $\mathcal{D=}\left\{ \mathbf{x}{}_{i},y_{i}\in\right\}_{i=1}^{N}$ – where $\mathbf{x}{}_{i}$ are $D$-dimensional vectors of covariates, $N$ is the total sample size and $y{}_{i}\mathbb{\in R}$ are response variables – the response variables are predicted using a potentially nonlinear regression model with additive Gaussian noise, i.e.: $y_{i}=f_{i}+\epsilon_{i}$ where $\epsilon_{i}\sim N(0{,\sigma}_{n}^{2})$. Inference then proceeds by placing a GP prior over this function then computing the posterior distribution using the canonical GPR predictive equations (17). This prior is uniquely specified by a mean ($m\left( x \right)$) and covariance ($k\left( x,x' \right)$) function. Here, without loss of generality we choose a mean function equal to zero and a generic covariance function combining linear and non-linear terms, i.e.:

$$k\left( \mathbf{x}_{i},\mathbf{x}_{j} \right)\mathbf{=}\mathbf{x}_{i}^{T}\mathbf{x}_{j}+\sigma_{f}\exp\left( -\frac{1}{2}\left( \mathbf{x}_{i}-\mathbf{x}_{j} \right)^{T}\boldsymbol{\Lambda}(\mathbf{x}_{i}-\mathbf{x}_{j}) \right)$$

Where $\sigma_{f}$ is a signal amplitude parameter for the nonlinear component and $\boldsymbol{\Lambda}$ is a diagonal matrix with $\mathcal{l}_{d}^{-2}$ along the leading diagonal. These are ‘automatic relevance determination’ parameters (17) that can down-weight irrelevant dimensions in the input space or emphasize important dimensions. Training a GP model refers to finding the optimal values for the model parameters which are: $\mathcal{l}_{1},\ldots,\mathcal{l}_{D},\sigma_{n}$ and $\sigma_{f}$. This is conveniently achieved by maximizing the logarithm of the model evidence (i.e. the denominator of Bayes rule). Finally, we compute a single subject Z-statistic image for each subject ($i$) and at each brain location ($j$) by computing:

$$\frac{y_{ij}-\hat{y}_{ij}}{\sqrt{\sigma_{ij}^{2}+\sigma_{nj}^{2}}}$$

Here, $\hat{y}_{ij}$is the predicted mean and predicted variance, $\sigma_{ij}$, which is combined with the true response ($y_{ij}$) and variance learned from the TD distribution ($\sigma_{nj}$). Because we estimate a separate noise parameter for each vertex, this should accommodate regional differences in population variation (for example, the estimated variance parameter will be higher in the regions where there is greater variation across individuals).

The Bayesian statistical model takes various sources of uncertainty into account, automatically making inferences more conservative in regions where data are sparse. Thus, the normative model can be estimated solely based on the NT cohort (N=233), and avoids enrichment for autism.

**Cross validation**

To assess generalizability, we used 10‐fold cross validation partitioning the data into 10 folds and repeatedly trained the model on 90% of the data, withholding the remaining 10% for estimating generalization performance. This was done 10 times so that each partition was excluded once. This is standard procedure in machine learning and provides unbiased estimates of the true generalizability.

**Structural and functional ROIs**

Structural ROIs were based on the Harvard-Oxford atlas (HOA (18)). The HOA was first coregistered using the nearest‐neighbor method to the symmetrical study‐specific template in MNI space and constrained to voxels in the study‐specific template. This resulted in 48 cortical and 8 subcortical right-hemisphere ROIs. Functional ROIs were based on reverse inference maps from the online meta‐analytic database neurosynth ([http://neurosynth.org](http://neurosynth.org/" \t "_blank), accessed June 2019) (19) using the search terms ‘language’, ‘motor’ (left-lateralized), ‘visuospatial’, ‘attention’ (right-lateralized) and ‘monitoring’, ‘mentalizing’ (no lateral bias). Maps were resliced to match the voxel resolution of the data, binarized, reflected along the *x* axis and the conjunction of right and left ROIs was used for the analyses to ensure symmetrical ROIs

**Detailed language delay results**

To assess the impact of LD, statistical second level-analyses were re-run by including LD (i.e., autism with LD, autism without LD, NT) as independent variable (instead of diagnostic group (i.e., autism and NT)). For follow-up analysis, we only selected those ROIs functionally related to language. For both extreme left- and rightward deviations, there was a significant main effect of LD (right: *F_(2)_*=10.5, *p*<0.001; left: *F_(2)_*=10.0, *p*<0.001); but not for sex (right: *F_(1)_*=0.1, *p*=0.8; left: *F_(1)_*=0.001, *p*=0.97), nor the sex-by-LD interactions (right: *F_(2)_*=1.5, *p*=0.22; left: *F_(2)_*=1.5, *p*=0.26) (see Figure 4). These results remained unchanged when controlling for FIQ and handedness (see Supplemental Information). For extreme rightward deviations, follow-up analyses showed that individuals with autism and LD were significantly different from both individuals with autism without LD (*t*_(213)_=2.5, *p*=0.01, *d*=0.32) and NT individuals (*t*_(152)_=4.6, *p*<0.001, *d*=0.58), while individuals with autism without LD did not reach a significant difference from NT individuals (*t*_(256)_=1.9, *p*=0.06, *d*=0.21). This stepwise pattern was overall more pronounced in males with autism than in females with autism. In contrast, for extreme leftward deviations, individuals with autism and LD were not different from individuals with autism without LD (*t*_(228)_=1.2, *p*=0.22, *d*=0.16), however both individuals with autism with and without LD were significantly different from NT individuals (NT vs. autism -LD: *t*_(182)_=4.4, *p*,0.001, *d*=0.53, NT vs. autism -noLD: *t_(_*_271)_=3.0, *p*=0.003, *d*=0.32). When matching individuals with autism with and without LD on symptom severity and age, results remained stable (see Figure S9).

When considering extreme deviations by structural ROIs related to language, there was a main effect of LD for extreme rightward deviations in the frontal operculum (*F_(2)_*=10.6, *p*<0.001) comprising parts of pars opercularis and the central operculum (*F_(2)_*=14.9, *p*<0.001). When considering deviations by functional ROIs, for extreme rightward deviations we detected a significant main effects of LD in the motor networks (*F_(2)_*=6.9, *p*=0.001). Rightward deviations in the language network were significant (*F_(2)_*=4.1, *p*=0.01), but did not survive FDR-correction. For all these, there was a stepwise pattern, with individuals with autism and LD showing more pronounced deviations.

**Controlling for confounds**

When controlling for FIQ and handedness in the second-level statistical analyses, results remained unchanged. However, note that due to limited availability of handedness information, this analysis was conducted in a smaller dataset (males with autism=93, females with autism=84, NT males=117, NT females=66). Overall, males and females with autism showed both significantly more extreme rightward deviations (*F_(1)_*=6.0, *p*=0.01) and leftward deviations (*F_(1)_*=7.7, *p*=0.006) compared to NT males and females. There were no sex differences (right: *F*=0.7, *p*=0.39; left: *F_(1)_*=0.02, *p*=0.9) and no sex-by-diagnosis interactions (right: *F_(1)_*=2.8, *p*=0.09; left: *F*=0.3, *p*=0.56). For both extreme left- and rightward deviations, there was a significant main effect of LD (right: *F_(2)_*=4.5, *p*=0.01; left: *F_(2)_*=7.3, *p*<0.001), however not for sex (right: *F_(1)_*=0.3, *p*=0.51; left: *F_(1)_*=0.3, *p*=0.56), or the sex-by-LD interactions (right: *F_(2)_*=1.5, *p*=0.23; left: *F_(2)_*=1.4, *p*=0.24).

**FDR-correction**

Overall, males and females with autism showed both significantly more extreme rightward deviations (*F_(1)_*=37.4, *p*<0.001) and leftward deviations (*F_(1)_*=28.4, *p*<0.001) compared to NT males and females. There were no sex differences (right: *F_(1)_*=1.5, *p*=0.22; left: *F_(1)_*=2.4, p=0.12) and no sex-by-diagnosis interactions for left (*F_(1)_*=2.8, *p*=0.1), however for extreme rightward deviations (*F_(1)_*=6.6, *p*=0.01), with females with autism showing the strongest rightward deviations. For both extreme left- and rightward deviations, there was a significant main effect of LD (right: *F_(2)_*=19.3, *p*<0.001; left: *F_(2)_*=10.6, *p*<0.001), and for sex for rightward deviations (*F_(1)_*=4.5, *p*=0.04), however not for sex for leftward deviations (*F_(1)_*=2.3, *p*=0.13). The sex-by-LD interaction was not significant for left (*F_(2)_*=2.2, *p*=0.11), however for rightward deviations (*F_(2)_*=3.5, *p*=0.03), showing that the stepwise pattern was more pronounced in males with autism than in females with autism. For extreme rightward deviations, follow-up analyses showed that individuals with autism and LD were not significantly different from each other (*t*_(227)_=0.8, *p*=0.43), however both individuals with autism with and without LD were significantly different from NT individuals (NT vs. autism-LD: *t*_(108)_=4.8, *p*<0.001), NT vs. autism-noLD: *t_(_*_169)_=3.8, *p*<0.001). For extreme leftward deviations, individuals with autism and LD were not different from individuals with autism without LD (*t*_(205)_=0.9, *p*=0.39), however both individuals with autism with and without LD were significantly different from NT individuals (NT vs. autism-LD: *t*_(136)_=4.6, *p*<0.001), NT vs. autism-noLD: *t_(_*_161)_=3.3, *p*=0.001).

**Exclusion of individuals with autism and intellectual disability (ID)**

We excluded individuals with autism and ID, resulting in a sample of 230 males with autism and 77 females with autism. Overall, males and females with autism showed both significantly more extreme rightward deviations (*F_(1)_*=8.8, *p*=0.003) and leftward deviations (*F_(1)_*=9.3, *p*=0.002) compared to NT males and females. There were no sex differences (right: *F_(1)_*=1.8, *p*=0.18; left *F_(1)_*=1.9, *p*=0.17) and no sex-by-diagnosis interactions (right: *F_(1)_*=0.6, *p*=0.42; left: *F_(1)_*<0.001, *p*=0.99). For both extreme left- and rightward deviations, there was a significant main effect of LD (right: *F_(2)_*=7.4, *p*=0.01; left: *F_(2)_*=9.0, *p*<0.001), however not for sex (right: *F_(1)_*=2.2, *p*=0.14; left: *F_(1)_*=0.8, *p*=0.36), or the sex-by-LD interactions (right: *F_(2)_*=0.3, *p*=0.73; left: *F_(2)_*=0.5, *p*=0.62).

**Matched sub-sample in autism (language-delayed vs. not-language-delayed)**

We used the python package ‘pymatch’ (<https://github.com/benmiroglio/pymatch>) to create age- and symptom-severity matched sub-samples of individuals with autism with and without language delay (LD). Matching resulted in a sample of 93 individuals with autism with LD (males=74; females=19) and 62 individuals with autism without LD (males=46; females=16). There were no significant differences between them on age (*t*_(126)_=0.1, *p*=0.91) and symptom severity as assessed by the CSS on the ADOS (*t*_(130)_=0.9, *p*=0.38). For both extreme left- and rightward deviations, there was a significant main effect of LD (right: *F_(2)_*=11.5, *p*<0.001; left: *F_(2)_*=10.5, *p*<0.001), however not for sex (right: *F_(1)_*=1.7, *p*=0.19; left: *F_(1)_*=0.04, *p*=0.83**.** The sex-by-LD interaction was not significant for left- (*F_(2)_*=1.6, *p*=0.2), however trending for rightward deviations (*F_(2)_*=3.0, *p*=0.05), showing that the stepwise pattern was more pronounced in males with autism than in females with autism (see Figure S9).

**Replication sample ABIDE I and II**

To assess replicability, we selected a sample from the publically available Autism Brain Imaging Data Exchange (ABIDE) I and II (20,21). Sites using the same scanner and acquisition parameters across ABIDE I and II were merged into one single site (KKI, NYU, OHSU, SDSU and UCLA 1) resulting in 15 sites in total. Selection criteria were as follows: a) subjects in the same age range as the primary analysis sample between 6-30 years of age, b) images with low image quality (due to excessive motion, insufficient coverage or brain abnormalities) and failed preprocessing were excluded; c) NT individuals with an ADHD diagnosis were excluded. This resulted in a sample of 418 males with autism, 95 females with autism, 473 NT males and 218 NT females. No information on language delay was available. For further details on demographic and clinical information see Table S2. T1 images were preprocessed with the same pipeline as describes in the methods of the main manuscript and as shown in Figure S2. No study-specific DARTEL template was created, but images were registered to the DARTEL template obtained from primary image analysis as described in the main text. Next, normative modeling and the creation of normative probability maps were done in accordance with the primary imaging analysis in the main text. Finally, we also tested for the spatial overlap between overlap maps in EU-AIMS LEAP (thresholded at 2%) and overlap maps in ABIDE (thresholded at 2%) to identify most strongly implicated regions across the datasets.

**Replicability results**

In line with the results of the EU-AIMS LEAP sample, males and females with autism of the combined ABIDE I and II dataset showed both significantly more extreme rightward deviations (*F_(1)_*=7.7, *p*=0.006, *d*=0.14) and leftward deviations (*F_(1)_*=17.1, *p*<0.001, *d*=0.24) compared to NT males and females (see Figure S10a-b). As in EU-AIMS LEAP, there were no sex differences (right: *F_(1)_*=1.1, *p*=0.3, *d*=0.14; left: *F_(1)_*=2.9, *p*=0.13, *d*=0.19) and no sex-by-diagnosis interactions (right: *F_(1)_*=3.6, *p*=0.06, *d*=0.26; left: *F_(1)_*=3.4, *p*=0.06, *d*=0.26). When considering deviations by functional ROIs that were significant in the primary analysis, individuals with autism also showed extreme rightward deviations in the motor network (*F_(1)_*=5.2, *p*=0.02, *d*=0.21), and extreme leftward deviations in the visuospatial network (*F_(1)_*=4.0, *p*=0.05, *d*=0.12). There were no significant group differences in the frontal operculum however (*F_(1)_*=0.2, *p*=0.6, *d*=0.02). Notably, regions with highest overlap in EU-AIMS LEAP and ABIDE for extreme rightward deviations were in the middle and superior temporal gyrus, precentral gyrus, hippocampus, putamen and intracalcerine cortex in males with autism and in the thalamus, parahippocampal gyrus, precuenus, intra- and supracalcerine cortex, superior frontal gyrus, frontal pole, orbitofrontal cortex and middle temporal gyrus in females with autism (see Figure 10a). Regions with highest overlap in EU-AIMS LEAP and ABIDE for extreme leftward deviations were in the thalamus and cerebellum in males with autism and in the caudate, anterior cingulate cortex, frontal pole and medial frontal cortex in females with autism (see Figure 10b).
