## Supplementary figures and images for "Atypical brain asymmetry in autism – a candidate for clinically meaningful stratification"

### Supplementary Figure 1

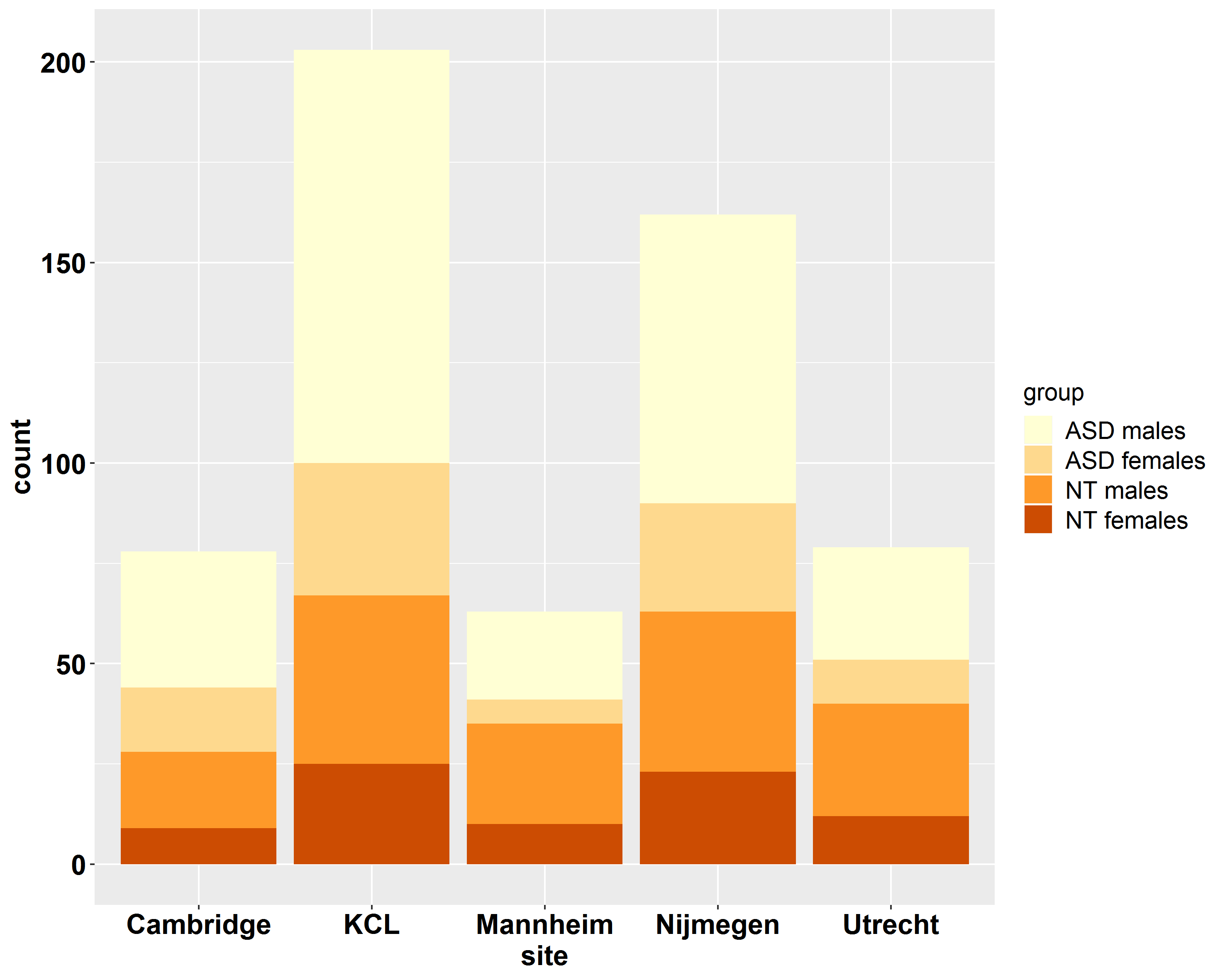

### Supplementary Figure 2

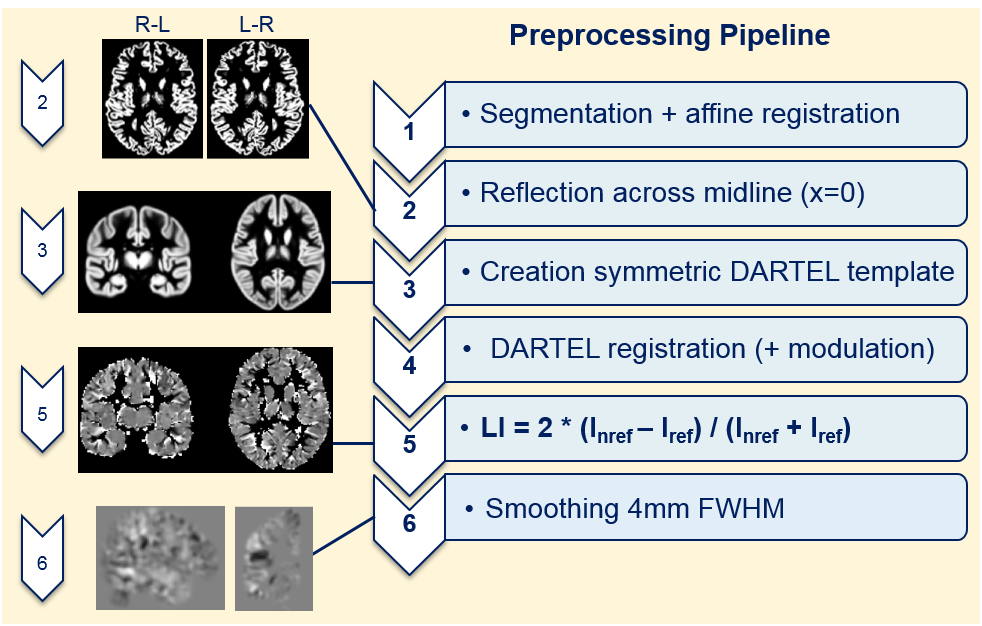

### Supplementary Figure 3

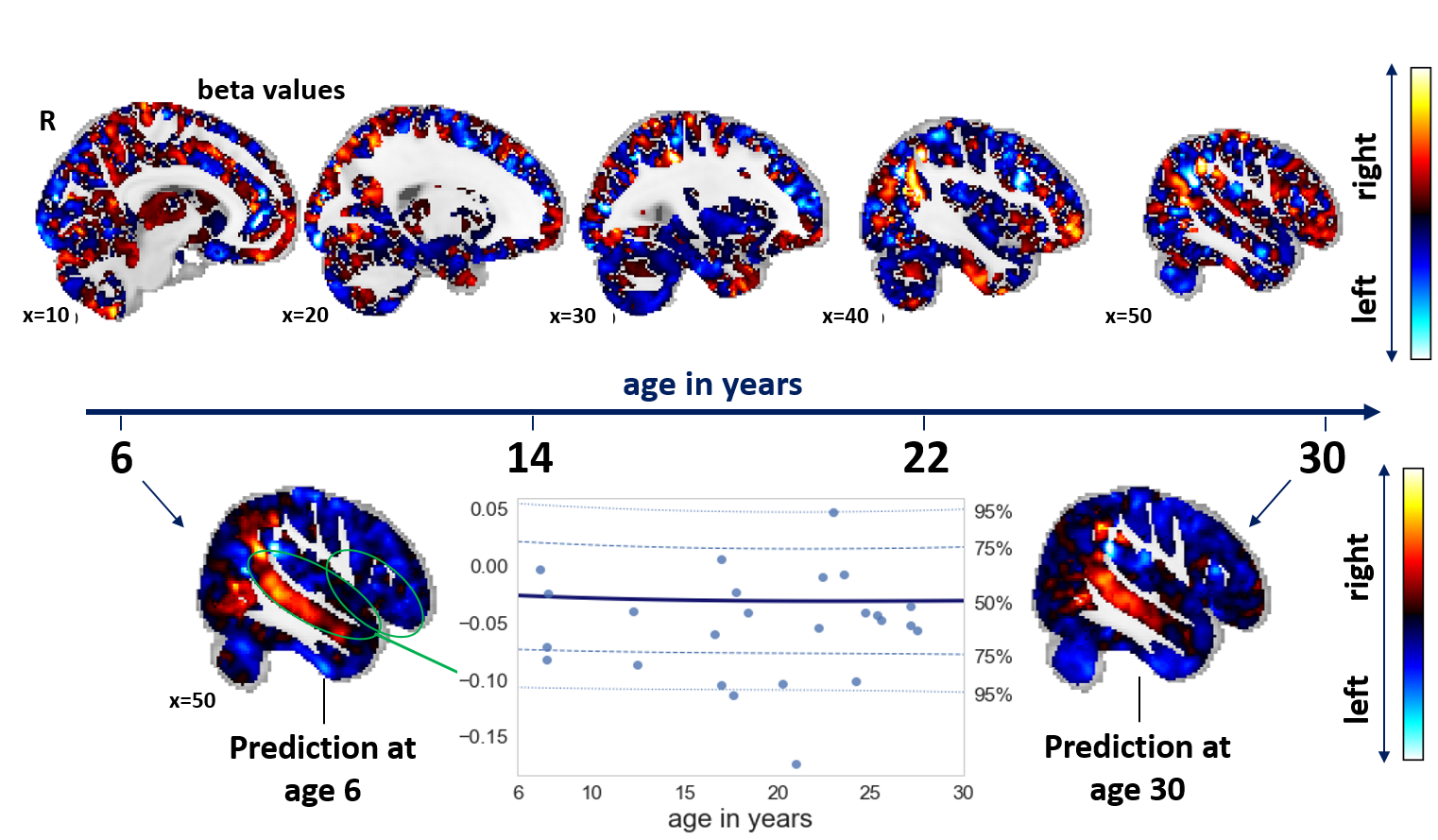

### Supplementary Figure 4a

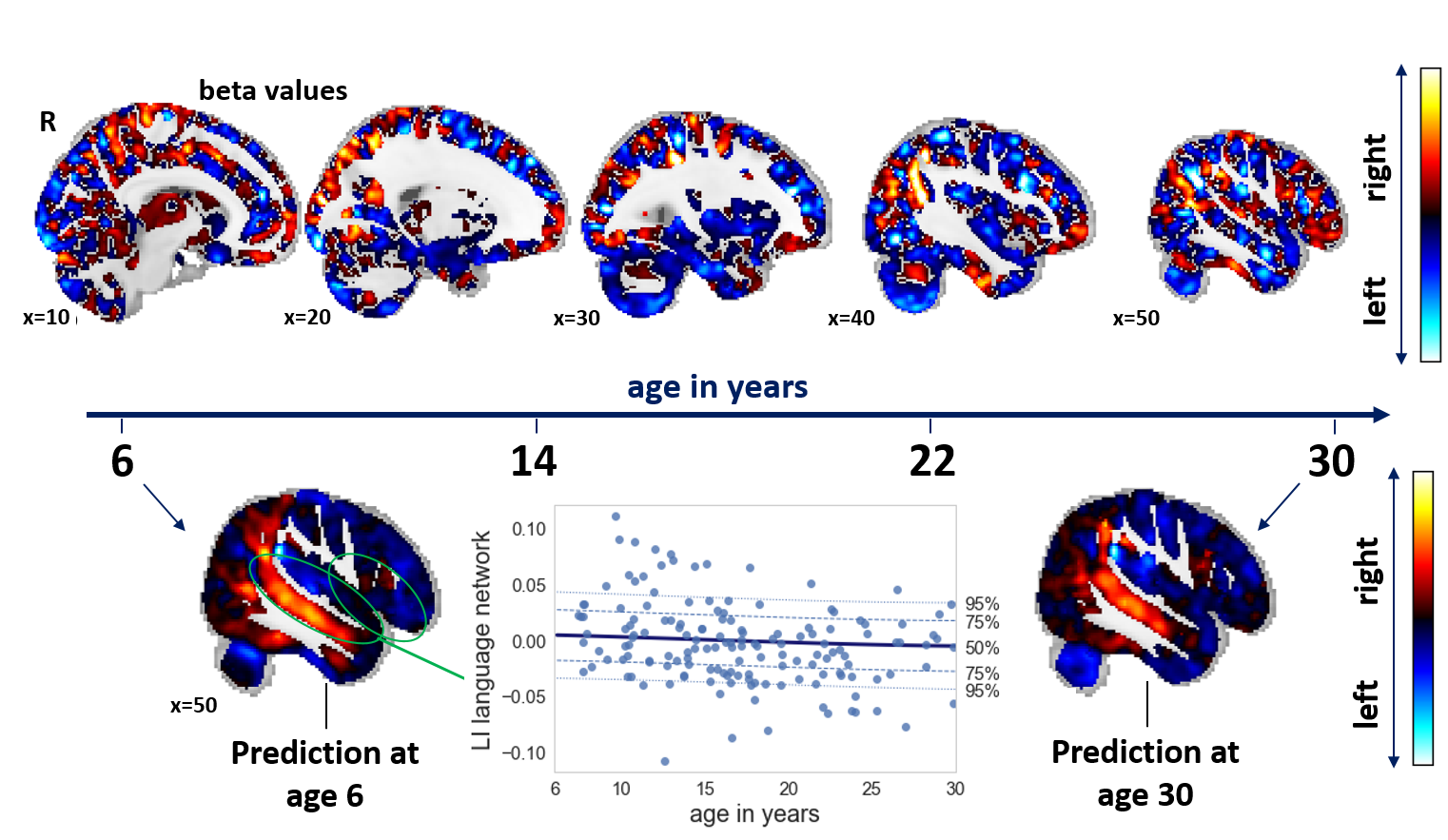

### Supplementary Figure 4b

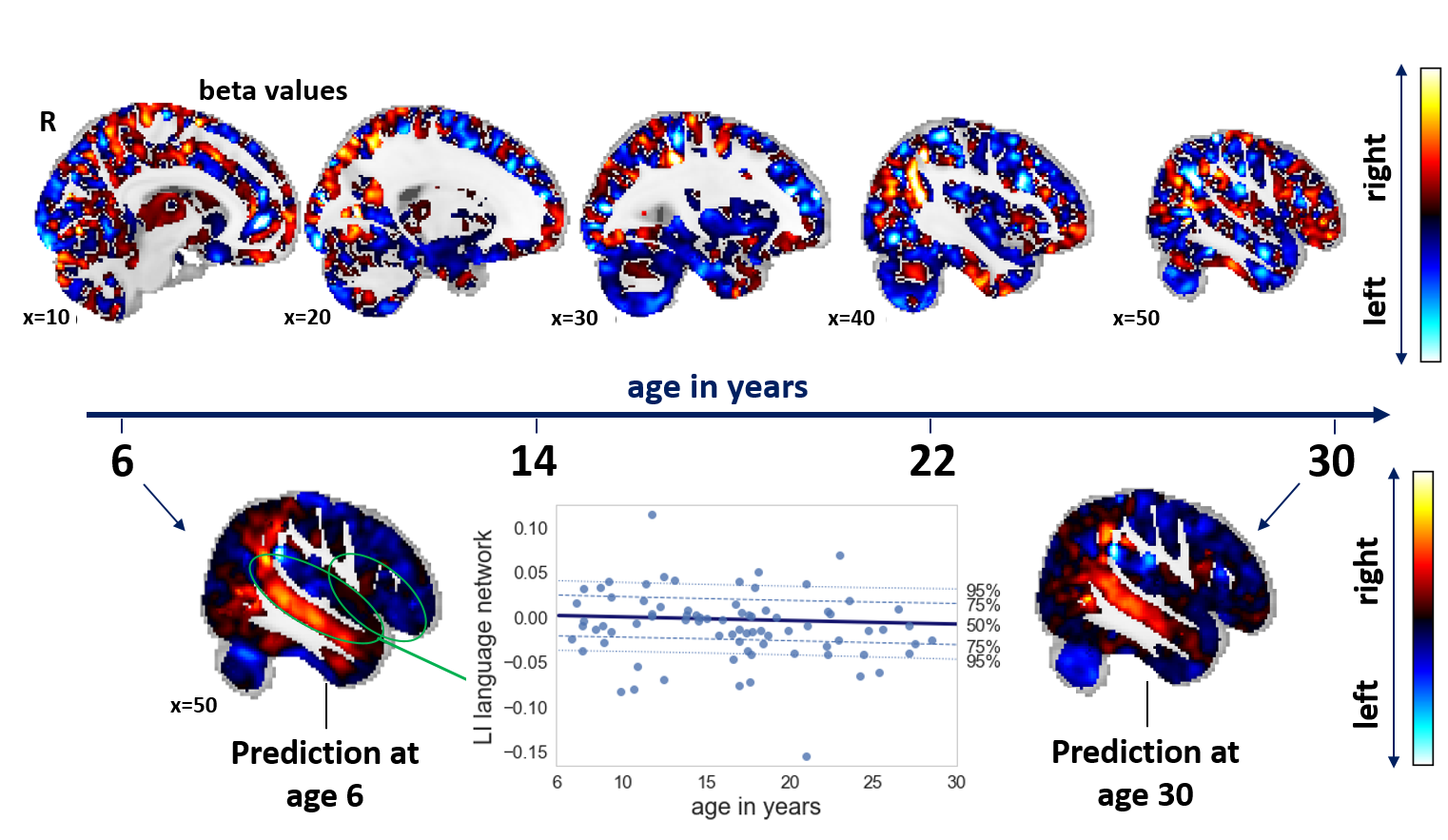

### Supplementary Figure 5

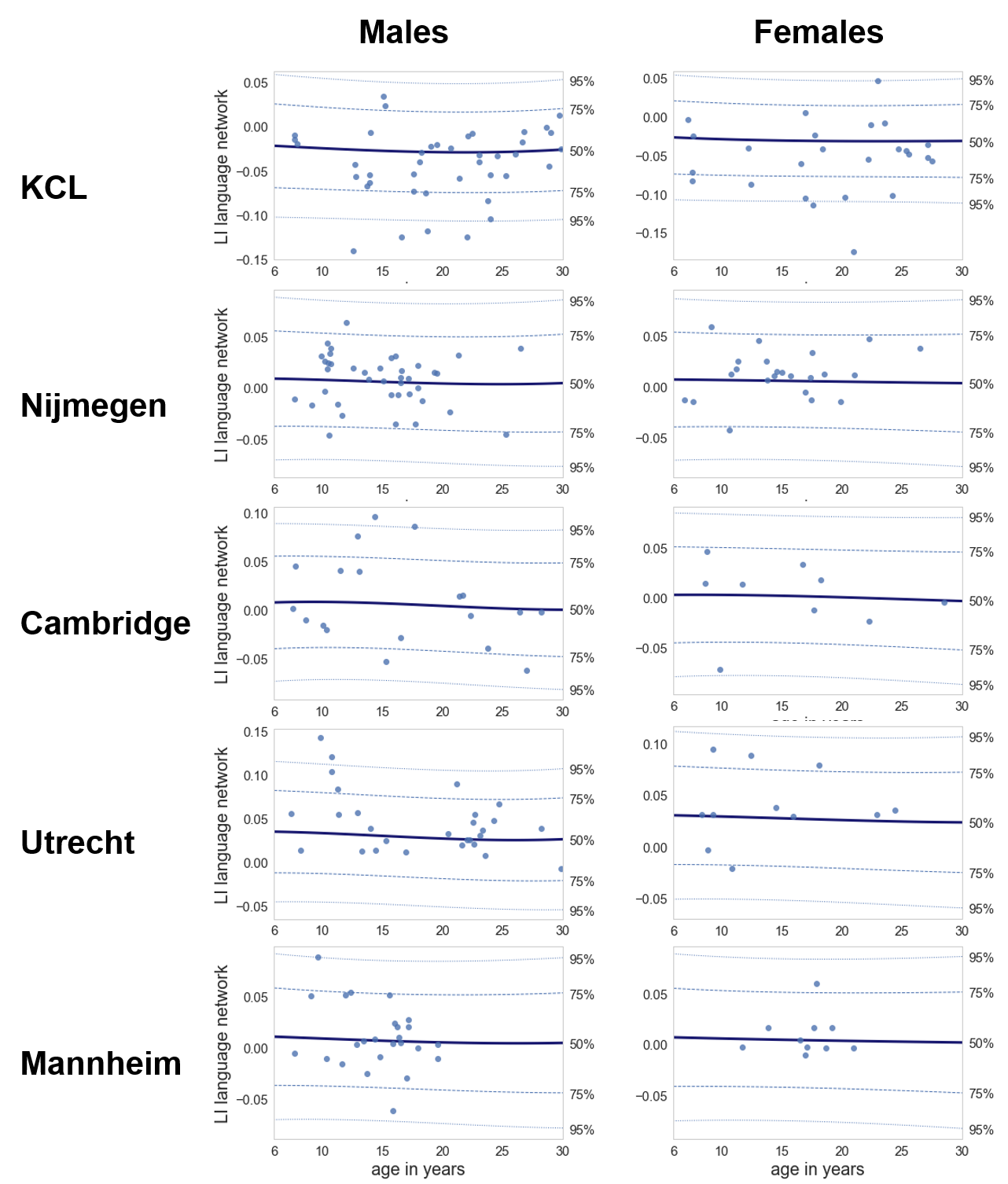

### Supplementary Figure 6

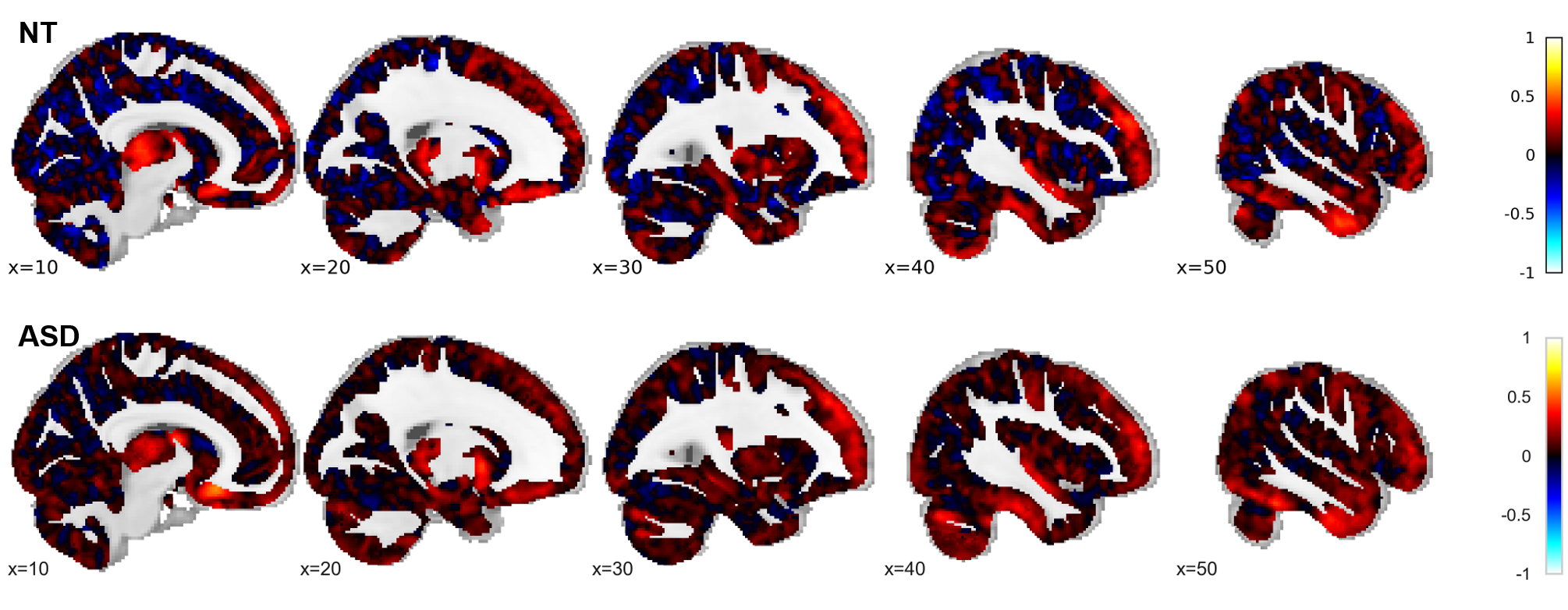

### Supplementary Figure 7

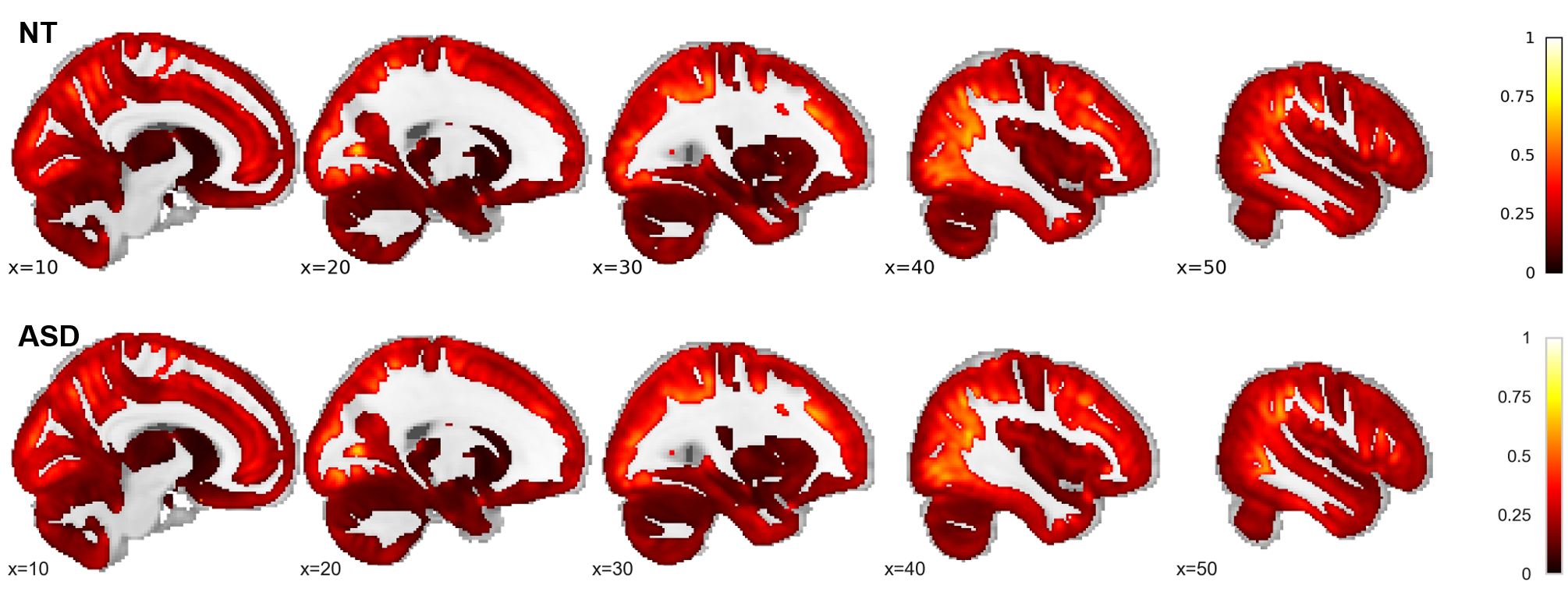

### Supplementary Figure 8

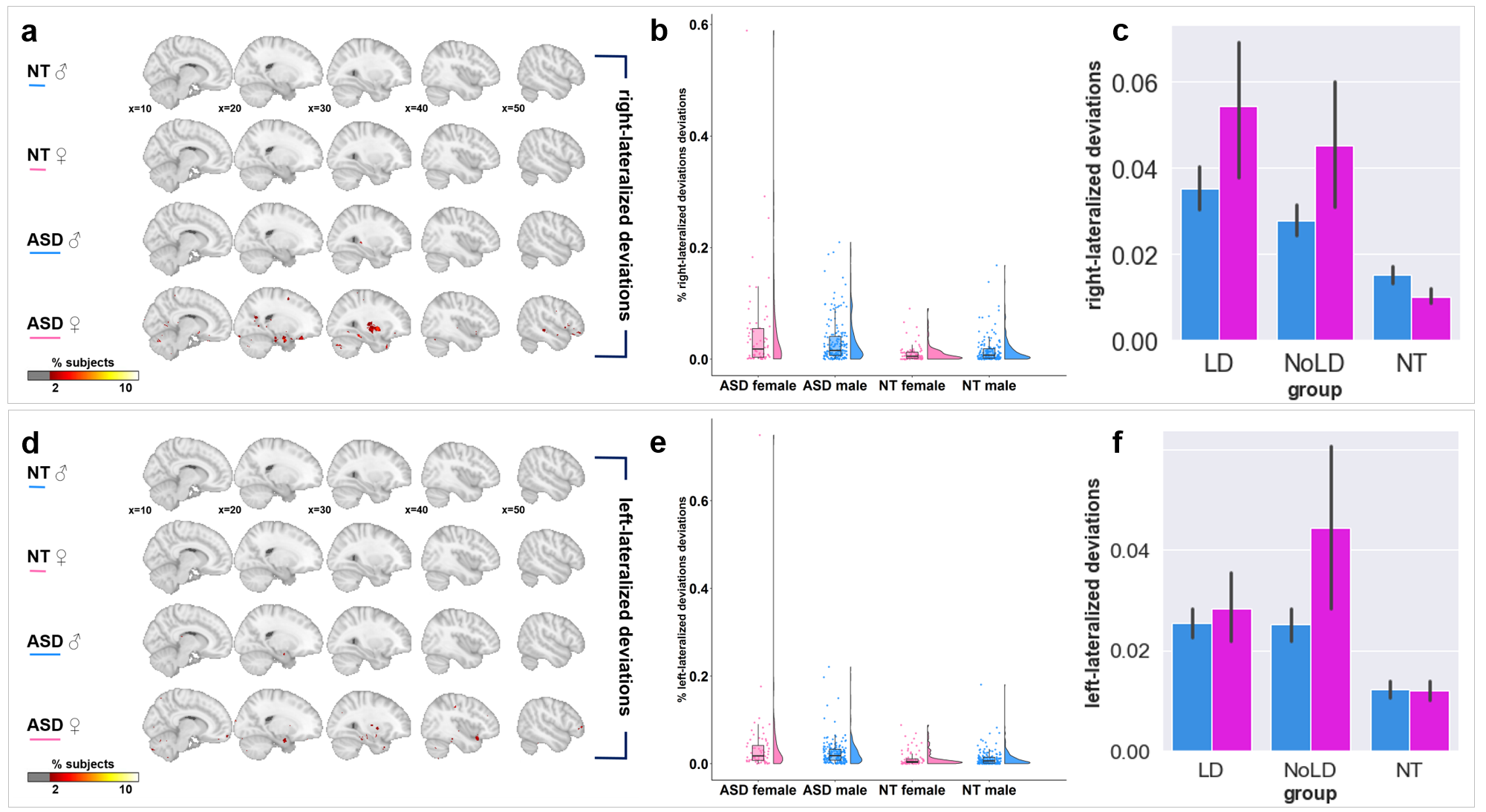

### Supplementary Figure 9

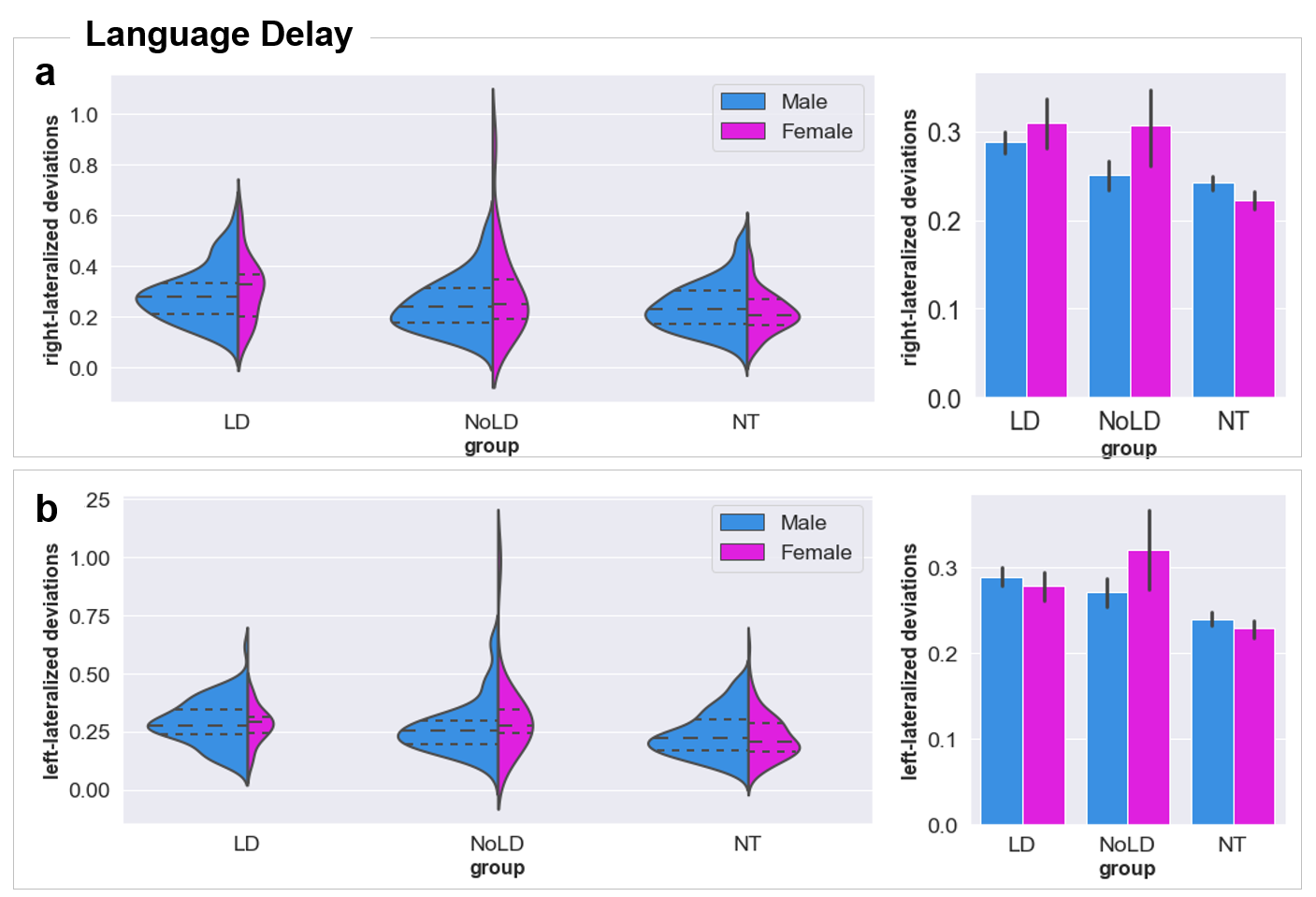

### Supplementary Figure 10

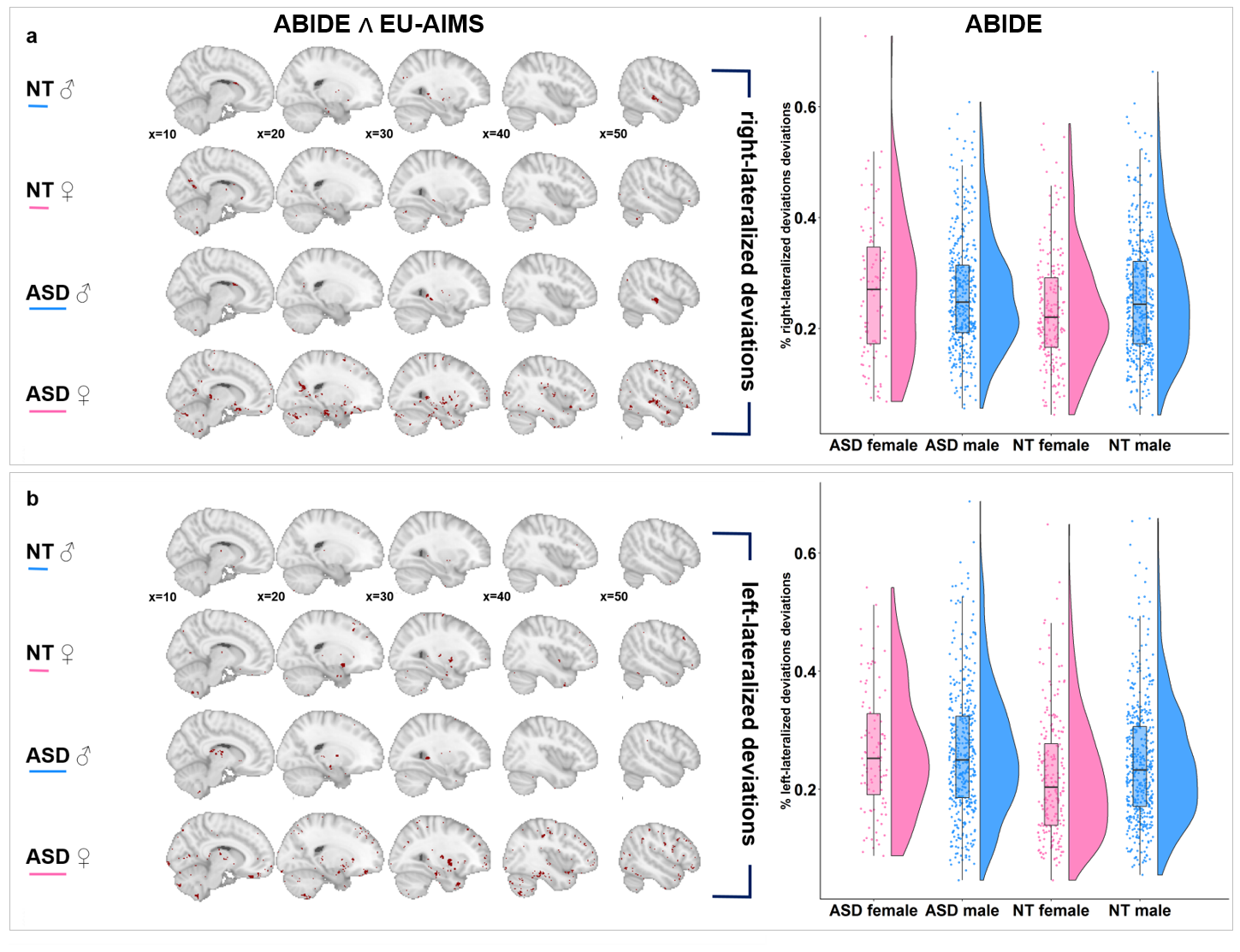
