## Supplementary material for "Atypical brain asymmetry in autism – a candidate for clinically meaningful stratification": Tables

**Table 1 – Demographic and clinical characterization of the LEAP sample**

|  | ASD M (N=259) | ASD F (N=93) | NT M (N=154) | NT F (N=79) |  |
| --- | --- | --- | --- | --- | --- |
|  | **Mean (SD)**  **[Range]** | **Mean (SD)**  **[Range]** | **Mean (SD)**  **[Range]** | **Mean (SD)**  **[Range]** | **Post-hoc** |
| Age | 16.8 (5.4)  [7.1-30.3] | 16.9 (6.1)  [6.8-30.3] | 17.1 (5.9)  [7.4-30.9] | 16.4 (5.8)  [6.9-28.5] | ns |
| Full-Scale IQ | 100 (18.9)  [56-148] | 97 (18.5)  [57-131] | 107 (15.1)  [53-142] | 107 (18.0)  [52-142] | (ASD M=ASD F) < (NT M=NT F) |
| Verbal IQ | 98 (18.8)  [52-160] | 97 (18.6)  [50-136] | 105 (16.3)  [46-142] | 107 (19.6)  [51-160] | (ASD M=ASD F) < (NT M=NT F) |
| Performance IQ | 101 (21.0)  [44-150] | 97 (19.7)  [55-133] | 108 (17.3)  [51-147] | 105 (18.4)  [58-139] | (ASD M=ASD F) < (NT M=NT F) |
| ADI-R |  |  |  |  |  |
| Social | 17.2 (6.4)  [1-28] | 15.5 (7.1)  [1-29] | - | - | ns |
| Communication | 13.8 (5.7)  [0-26] | 12.4 (5.3)  [0-24] | - | - | ASD M>ASD F |
| RRB | 4.5 (2.7)  [0-12] | 3.8 (2.6)  [0-10] | - | - | ASD M>ASD F |
| ADOS-2 |  |  |  |  |  |
| Social-Affect | 6.2 (2.6)  [1-10] | 5.3 (2.5)  [1-10] | - | - | ASD M>ASD F |
| RRB | 4.9 (2.8)  [1-10] | 4.3 (2.6)  [1-9] | - | - | ns |
| CSS total | 5.6 (2.8)  [1-10] | 4.5 (2.5)  [1-10] | - | - | ASD M>ASD F |
| Handedness | 167 (R)  34 (L)  7 (A) | 71 (R)  9 (L)  4 (A) | 105 (R)  9 (L)  3 (A) | 56 (R)  8 (L)  2 (A) | (ASD M=ASD F) < (NT M=NT F) |
| ADHD | 106 (ADHD+)  109 (ADHD –) | 34 (ADHD+)  52 (ADHD –) | - | - | ns |
| LD | 76 (LD)  100 (NoLD) | 19 (LD)  47 (NoLD) | - | - | ns |

Abbreviations: ASD=Autism Spectrum Disorder; NT=neurotypical, M=males, F=females, ADI-R= Autism Diagnostic Interview-Revised, ADOS=Autism Diagnostic Observation Schedule, RRB=restricted, repetitive behavior, CSS=calibrated severity score, ADHD= attention-deficit hyper-activity disorder, LD=language delay.

**Supplementary Table 1 – Summary of acquisition parameters across sites**

| Site | Manufacturer | Model | Software Version | Acquisition sequence | Coverage | Slices | Thickness [mm] | Resolution [mm^3^] | TR [s] | TE [ms] | FA [°] | FOV |
| --- | --- | --- | --- | --- | --- | --- | --- | --- | --- | --- | --- | --- |
| Cambridge | Siemens | Verio | Syngo MR B17 | Tfl3d1_ns | 256*256 | 176 | 1.2 | 1.1*1.1*1.2 | 2.3 | 2.95 | 9 | 270 |
| London | GE Medical systems | Discovery mr750 | LX MR DV23.1_V02_1317.c | SAG ADNI GO ACC SPGR | 256*256 | 196 | 1.2 | 1.1*1.1*1.2 | 7.31 | 3.02 | 11 | 270 |
| Mannheim | Siemens | TimTrio | Syngo MR B17 | MPRAGE ADNI | 256*256 | 176 | 1.2 | 1.1*1.1*1.2 | 2.3 | 2.93 | 9 | 270 |
| Nijmegen | Siemens | Skyra | Syngo MRD13 | Tfl3d1_16ns | 256*256 | 176 | 1.2 | 1.1*1.1*1.2 | 2.3 | 2.93 | 9 | 270 |
| Rome | GE Medical systems | Signa HDxt | 24/LX/MR HD16.0_V02_1131.a | SAG ADNI GO ACC SPGR | 256*256 | 172 | 1.2 | 1.1*1.1*1.2 | 5.96 | 1.76 | 11 | 270 |
| Utrecht | Philips Medical Systems | Achieva/  Ingenia CX | 3.2.3, 3.2.3.1 | ADNI GO 2 | 256*256 | 170 | 1.2 | 1.1*1.1*1.2 | 6.76 | 3.1 | 9 | 270 |

**Supplementary Table 2 - Demographic and clinical characterization of the ABIDE sample**

|  | ASD M (N=418) | ASD F (N=95) | NT M (N=473) | NT F (N=218) |  |
| --- | --- | --- | --- | --- | --- |
|  | **Mean (SD)**  **[Range]** | **Mean (SD)**  **[Range]** | **Mean (SD)**  **[Range]** | **Mean (SD)**  **[Range]** | **Post-hoc** |
| Age | 12.8 (4.1)  [6.8-30.0] | 12.6 (4.3)  [6.9-27.0] | 12.7 (4.2)  [7.1-29.0] | 12.5 (4.7)  [7.0-29.9] | ns |
| Full-Scale IQ^a^ | 106 (17.6)  [49-149] | 104 (17.8)  [66-147] | 113 (12.6)  [71-148] | 113 (13.0)  [80-149] | (ASD M=ASD F) < (NT M=NT F) |
| Verbal IQ^b^ | 107 (18.7)  [45-180] | 104 (17.7)  [62-145] | 114 (13.2)  [73-147] | 113 (14.6)  [83-156] | (ASD M=ASD F) < (NT M=NT F) |
| Performance IQ^c^ | 106 (17.3)  [59-149] | 102 (18.1)  [53-148] | 109 (13.4)  [62-147] | 109 (13.3)  [79-145] | (ASD M=ASD F) < (NT M=NT F) |
| ADI-R |  |  |  |  |  |
| Social^d^ | 19.6 (5.4)  [4-30] | 19.0 (6.4)  [0-30] | - | - | ns |
| Communication^e^ | 15.7 (4.6)  [2-25] | 14.8 (5.3)  [0-25] | - | - | ns |
| RRB^e^ | 6.0 (2.5)  [0-13] | 5.7 (2.5)  [0-12] | - | - | ns |
| ADOS-2 |  |  |  |  |  |
| Social-Affect^f^ | 9.2 (3.8)  [2-20] | 9.1 (3.5)  [4-18] | - | - | ns |
| RRB^g^ | 2.9 (1.9)  [0-8] | 2.6 (1.5)  [0-5] | - | - | ns |
| CSS total^h^ | 6.9 (2.2)  [1-10] | 6.8 (1.9)  [2-10] | - | - | ns |
| Handedness | 171 (R)  14 (L)  16 (A) | 43 (R)  6 (L)  3 (A) | 207 (R)  14 (L)  14 (A) | 128 (R)  5 (L)  4 (A) | (ASD M=ASD F) < (NT M>NT F) |

Abbreviations: ASD=Autism Spectrum Disorder; NT=neurotypical, M=males, F=females, ADI-R= Autism Diagnostic Interview-Revised, ADOS=Autism Diagnostic Observation Schedule, RRB=restricted, repetitive behavior, CSS=calibrated severity score ^a^Full-Scale IQ information was available for 384 ASD M, 87 ASD F, 435 NT M and 197 NT F. ^b^Verbal IQ information was available for 330 ASD M, 76 ASD F, 382 NT M and 159 NT F. ^c^Performance IQ information was available for 335 ASD M, 78 ASD F, 398 NT M and 173 NT F. ^d^ADI-R Social Scores were available for 341 ASD M and 74 ASD F. ^e^ADI-R Communication and RRB Scores were available for 342 ASD M and 74 ASD F. ^f^ADOS-Gotham Social-Affect Scores were available for 132 ASD M and 28 ASD F. ^g^ADOS-Gotham RRB Scores were available for 134 ASD M and 29 ASD F. ^h^ADOS-Gotham Severity Scores were available for 134 ASD M and 30 ASD F.
